# Constrained Evolutionary Funnels Shape Viral Immune Escape

**DOI:** 10.1101/2025.10.26.684604

**Authors:** Marian Huot, Dianzhuo Wang, Eugene Shakhnovich, Rémi Monasson, Simona Cocco

**Affiliations:** Laboratory of Physics of the École Normale Supérieure, CNRS UMR 8023 and PSL Research, Sorbonne Université, 24 rue Lhomond, Paris, France; Department of Chemistry and Chemical Biology, Harvard University, Cambridge, MA; John A. Paulson School of Engineering and Applied Sciences, Harvard University, Cambridge, MA

**Keywords:** Viral adaptation, Mutational pathways, Restricted Boltzmann machines, SARS-CoV-2, Antibody escape, Protein evolution

## Abstract

Understanding how viral proteins adapt under immune pressure while preserving viability is crucial for anticipating antibody-resistant variants. We present a probabilistic framework that predicts viral escape trajectories and shows that immune evasion is channeled into a small set of viable “escape funnels” within the vast mutational space. These escape funnels arise from the combined constraints of protein viability and antibody escape, modeled using a generative model trained on homologs and deep mutational scanning data. We derive a mean-field approximation of evolutionary path ensembles, enabling us to quantify both the fitness and entropy of escape routes. Applied to SARS-CoV-2 receptor binding domain, our framework reveals convergent evolution patterns, predicts mutation sites in variants of concern, and explains differences in antibody-cocktail effectiveness. In particular, cocktails with de-correlated escape profiles slow viral adaptation by forcing longer, higher-cost escape paths.

**SIGNIFICANCE:** Viruses evolve to evade our immune defenses, but with constraints. Like navigating a minefield, each step toward immune escape comes at the potential cost of structural stability and functionality. We show that despite the vast mutational space, immune escape is funneled into a small set of predictable pathways. Using a statistical-physics model grounded in antibody experiments and SARS-CoV-2 epidemiology data, we identify these escape funnels—enabling therapies designed to block them before they are ever used.

## INTRODUCTION

Anticipating the evolutionary trajectories of viruses is pivotal to pandemic preparedness, driven by their continuous adaptation under immune pressure^1,2^. Immune responses elicited by prior infections and vaccinations impose selective pressure that significantly shapes viral evolution, resulting in the emergence of variants characterized by mutations that enhance transmissibility and enable antibody evasion^3–5^.

Several studies^6–8^ documented striking cases of convergent evolution in SARS-CoV-2, showing that distinct viral lineages can independently acquire similar mutations in the receptor binding domain (RBD). These convergent mutations cluster at key residues, enabling antibody escape while preserving binding to cell receptor ACE2. Cao et al.^6^ further demonstrated that this pattern is especially evident in Omicron subvariants, where immune imprinting from earlier exposures narrows the diversity of neutralizing antibodies, intensifying selective pressures and driving convergent evolutionary pathways.

Significant progress has been made in demonstrating the effects of individual viral mutations, largely through powerful experimental approaches such as deep mutational scanning (DMS)^6,9–17^. These techniques provide invaluable data on how single amino acid mutations in the SARS-CoV-2 spike protein or RBD influence key viral traits, including receptor-binding affinity, viral entry, and antibody neutralization. Such insights have proven critical for viral surveillance and have even enabled the forecasting of evolutionary success for specific viral clades^18–21^. Yet, quantifying how these mutations combine and interact across entire evolutionary trajectories remains a largely unexplored challenge. While many computational approaches can now predict the fitness effects of individual mutations^22–24^ or low-order combinations^25^, they typically overlook the collective, high-dimensional epistatic constraints that ultimately govern which mutational paths are viable. In realistic protein fitness landscapes, epistatic interactions emerge progressively as mutations accumulate, dynamically reshaping the accessibility of subsequent mutations and giving rise to long, emergent evolutionary timescales^26^.

Recent theoretical advances, including transition path sampling^27,28^ and navigability in highdimensional genotype–phenotype maps^29^, have begun to reveal how epistasis and evolutionary constraints jointly shape accessible evolutionary paths. Yet, a comprehensive, quantitative framework for predicting how functional and immune selection co-constrain the ensemble of escape trajectories, validated against real-world data, remains lacking.

Here, we present a probabilistic framework that characterizes immune escape as a constrained dynamical process through sequence space. By jointly modeling protein viability and antibody evasion, we demonstrate that viral adaptation is funneled through a remarkably small number of mutational trajectories, which we term “escape funnels”. To quantify these funnels, we construct a fitness landscape where viability is captured by Restricted Boltzmann Machines (RBMs). Simultaneously, immune escape is modeled using antibody-specific binding scores derived from deep mutational scanning (DMS).

We benchmark this approach on a solvable lattice protein model before applying it to the SARS-CoV-2 RBD. Crucially, we derive a mean-field approximation of the path ensemble, which enables the tractable computation of the free energy, entropy, and continuity of escape trajectories. This analytical approach reveals a sharply reduced set of viable escape paths, offers an explanation for the convergent evolution observed in real-world variants, and quantifies how specific antibody combinations reshape the mutational landscape to constrain viral evolution.

## RESULTS

### Escape paths

To investigate how proteins can escape immune pressure while maintaining viability, we developed a framework to sample evolutionary paths^27,28^ under immune pressure (Figure 1). Each path is defined as a sequence of amino acid variants 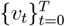, starting from the wildtype *v*_0_. The overall path probability is factorized as

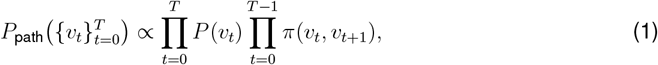

where the transition factor *π*(*v*_*t*_, *v*_*t*+1_) increases with the similarity between the sequences *v*_*t*_, *v*_*t*+1_. In practice, we use *π*(*v*_*t*_, *v*_*t*+1_) = **1** {*d*_*H*_(*v*_*t*_, *v*_*t*+1_)≤1} such that paths are constructed through single-residue mutations per step. Each *P* (*v*_*t*_) balances two competing pressures: the requirement to maintain protein viability (which includes stability and capacity to bind to ACE2 receptor) and the need to escape antibody binding.

**Fig. 1:**
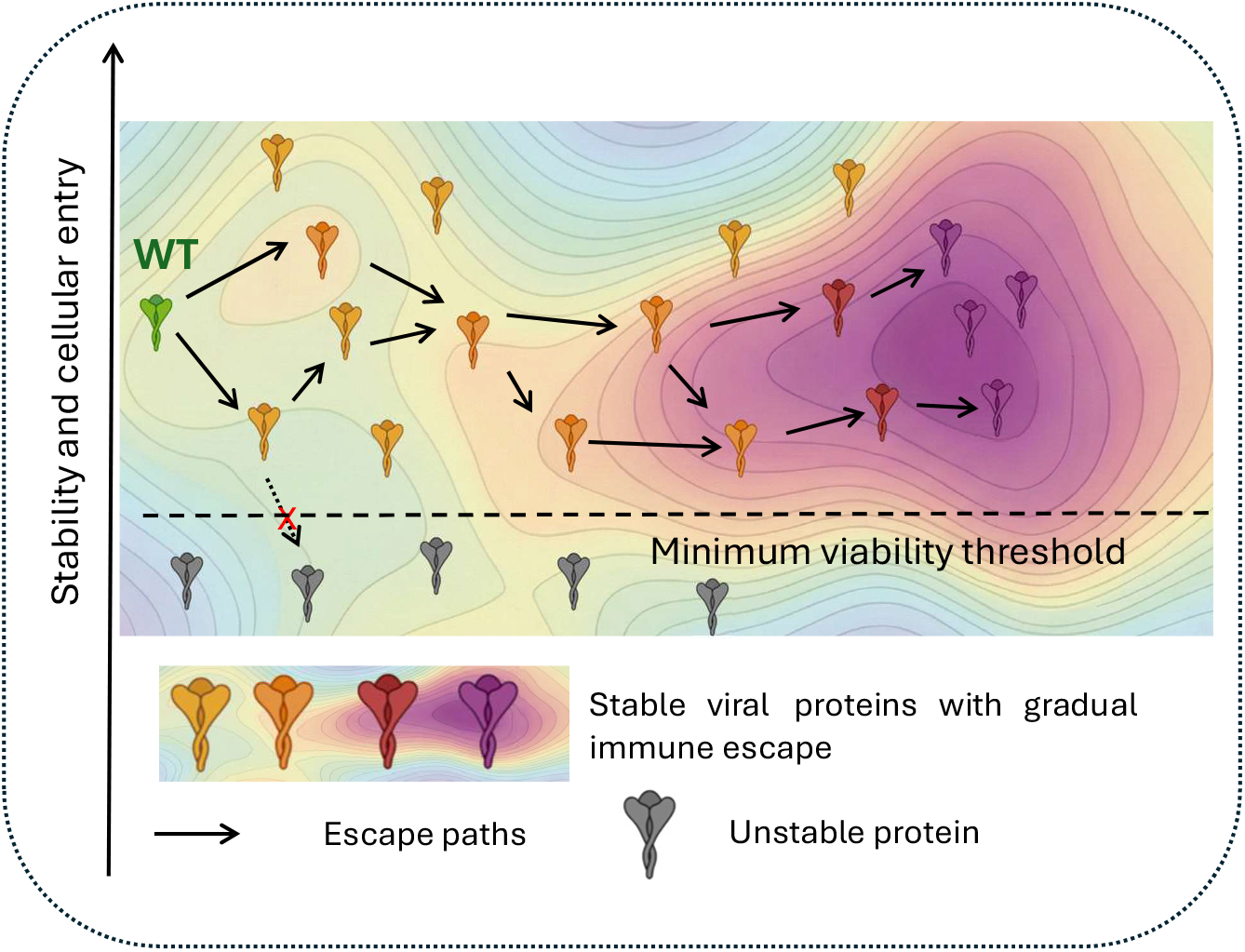
Evolutionary paths toward antibody escape. Schematic representation of plausible evolutionary trajectories from the wildtype to antibody-escaping variants. Paths progress through a constrained sequence space, funneling through a small set of viable intermediates that gradually accumulate immune escape mutations.

To model protein viability, we learn the probability of each protein sequence *v* = (*v*_1_, …, *v*_*N*_ ) using a Restricted Boltzmann Machine (RBM)^30,31^. The RBM is trained on a multiple sequence alignment of protein variants and serves as a generative model for viable sequences. Differently from previous works^32^ the RBM is trained on artificial multiple sequence alignment using ESM-inverse Folding, conditioned to the RBD-ACE2 complex structure (Methods).The RBM captures both local constraints on individual sites (through site-specific fields *g*_*i*_) and higher-order epistatic interactions (through a hidden layer). In this model, each visible unit *i* represents the amino-acid identity *v*_*i*_ at site *i* in the sequence, while each hidden (latent) unit *μ* summarizes a collective interaction pattern across sites via the input *I*_*μ*_(*v*) =Σ_*i*_ *w*_*iμ*_(*v*_*i*_), where *w*_*iμ*_ encodes the contribution of residue *v*_*i*_ at site *i* to hidden unit *μ*. The probability of observing a sequence is then:

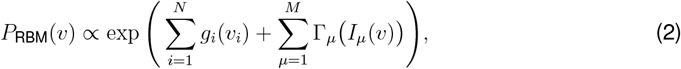

where the non-linear function Γ_*μ*_ is the cumulant generating function associated with hidden unit *μ*.

To model immune escape, we define

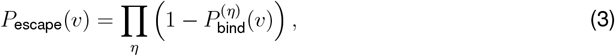

where 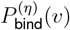 denotes the binding probability of sequence *v* to antibody *η*, and the product runs over a pool of antibodies.

The full fitness model

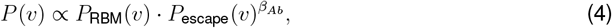

combines these two contributions and determines the probability of visiting each sequence along the evolutionary path. *β*_*Ab*_ enables to control escape and is a proxy for immune pressure.

To explore such trajectories, we extended a Monte Carlo sampling algorithm of transition paths (MCMC) previously introduced^27^ (see Methods). Each path is constructed under adjacency constraints (single-residue mutations per step), ensuring evolutionary continuity while respecting the underlying sequence probability in (4).

### Lattice protein model

We first benchmark this approach in a controlled setting, and apply it to lattice proteins (LP)^33,34^, a simplified yet physically grounded model of protein folding (Figure 2a). LP sequences consist of 27 amino acids folding into compact, self-avoiding walks on a 3× 3×3 cubic lattice, which defines the possible structures *S*. In this model, the ground-truth viability of a sequence *V* is the probability *P*_nat_(*S*_nat_ |**v**) that it folds in the native structure *S*_nat_ rather than in one of the many competing structures (see Methods). To mimic the situation in which we do not know the groundtruth fitness, we use the RBM trained on a set of high-*P*_nat_ LP sequences, as a surrogate model for *P*^nat 35^.

**Fig. 2:**
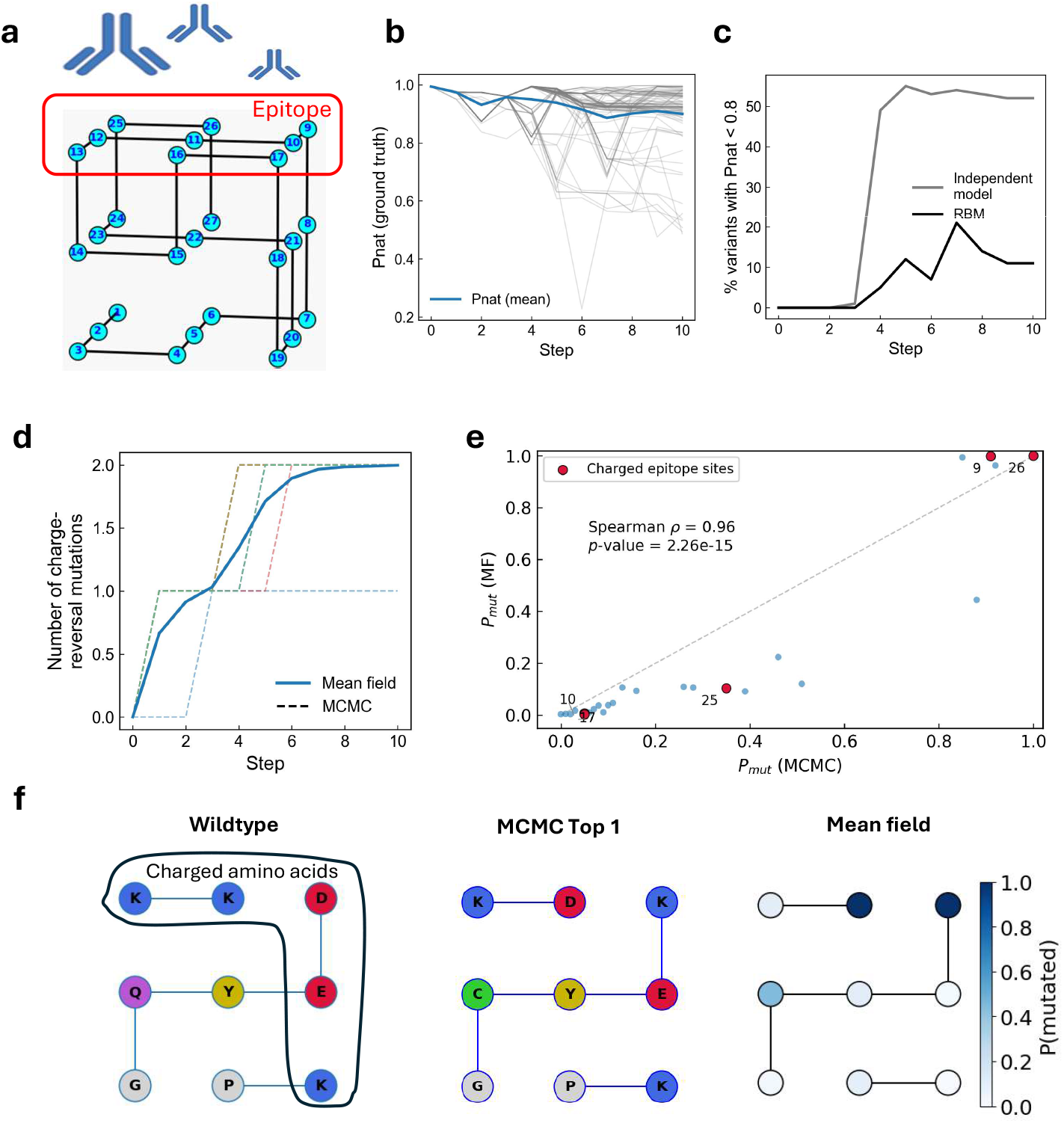
Folding constraints shape convergent escape routes under immune pressure on lattice protein model. **(a)** Native fold structure *S*_nat_, with the antibody-targeted epitope (upper face) highlighted in red. During escape, the protein must accumulate charge-reversal epitope mutations while preserving high folding probability. **(b)** Evolution of folding probability *P*_nat_ along sampled escape paths. The blue line indicates the mean across paths; grey lines show individual trajectories. **(c)** Comparison of the percentage of proteins with low folding probability in evolutionary paths generated by the RBM and a site-independent model. The lack of epistasis in the site-independent model leads to a rapid decrease in the number of viable variants after 4 mutations. **(d)** Accumulation of epitope charge-reversal mutations along escape paths. Each dashed line represents a single MCMC path, while blue line represents mean field trajectory. **(e)** Comparison site-wise probability of being mutated at the final step of the mean-field trajectory and in MCMC trajectories. **(f)** Comparison of wildtype epitope, most representative epitope variant at the terminal step of MCMC-sampled escape paths and site-wise epitope probability of being mutated at the final step of the mean-field trajectory. Amino-acid colors denote biochemical classes (positively charged in blue, negatively charged in red).

To represent the immune pressure exerted by an antibody, we introduce a binding probability that decays upon charge-reversing substitutions within a targeted epitope:

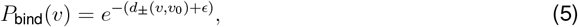

where *d*_*±*_(*v, v*_0_) counts the epitope mutations of opposite charge between candidate *v* and wildtype *v*_0_, and acts as binding energy. The term *ϵ* ≪ 1 is a regularization term as it ensures a finite log-escape probability, see Eq. (3), even for the unmutated wildtype.

We then sample escape paths of length *T* = 10 using our MCMC algorithm, starting from an initial sequence with high *P*_nat_ (Figure S1). Choosing *T* = 10 allows for a sequence divergence of ∼ 37%, comparable to the natural variability observed in the HCV E2 glycoprotein and HIV1 envelope protein^36,37^. Despite immune pressure, the sampled paths maintain high folding probabilities throughout the trajectory (Figure 2b): while individual paths exhibit some variability, the mean folding probability remains as high as wildtype folding probability, demonstrating that the RBM constraint successfully preserves structural integrity under immune-driven evolution. In particular, when using an independent model instead of the RBM, we find a high number of variants with lower *P*_nat_ after a few mutations (Figure 2c), showing the RBM’s ability to capture epistatic interactions between sites matters.

### Mean field for path sampling

A key advantage of using an RBM for sequence modeling is that its latent-unit structure admits a tractable mean-field (MF) approximation of the path ensemble, providing theoretical insight and scaling beyond sampled trajectories^27^. In the MF framework of statistical physics, each path is characterized via three sets of order parameters describing path sequences *v*_*t*_: (1) the sequence dependent inputs to each of the RBM hidden units 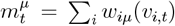, (2) the binding energy per antibody 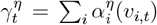 and (3) a continuity parameter along the path 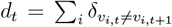 per antibody encoding the hamming distance between successive sequences (see Methods).

These parameters summarize the trajectory at each time step *t*, and the probability of sampling paths with such parameters is approximated as a Boltzmann weight

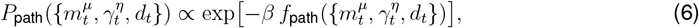

where the path free energy *f*_path_ can be exactly computed (see Methods). Here, *β* plays the role of an inverse temperature, allowing us to select paths of variable quality.

To validate the framework, we applied the MF formulation to our lattice model. We found that the MF model recovers the escape dynamics, specifically the occurrence of two charge-reversal mutations observed in MCMC paths (Figure 2d). Additionally, the MF model correctly predicts the specific mutated sites. It reveals that while many routes are possible given the five charged epitope residues, the system favors sites 9 and 26 (Figures 2e); this selectivity allows for an alternating arrangement of positive and negative charges on the epitope (Figures 2f), enabling escape without compromising *P*_nat_. Finally, the MF path closely matches the geometry of individual MCMC trajectories when projected onto the order-parameter space (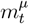, *γ*_*t*_) (Figure S2).

### Application to SARS-CoV-2 receptor binding domain

We next apply our framework to a more realistic setting by modeling SARS-CoV-2 receptor binding domain (RBD) sequences under functional and immune constraints. Viability and binding to ACE2 are modeled using an RBM trained on 1000 synthetic homologs generated with ESM-Inverse Folding, conditioned on the RBD–ACE2 complex structure. Antibody escape is incorporated using experimental deep mutational scanning (DMS) data^32,38^, for a fixed set of 29 antibodies identified early in the pandemic ^15^ (see Methods). Although additional experimental DMS datasets became available later^6^, this panel spans all four antibody classes and offers high-quality coverage with few missing single mutants. The objective is to determine whether an escape path exists: a rapid route with few mutational steps that circumvents the specified antibody set while maintaining protein viability. Unlike previous approaches^19,32^, we do not explicitly model antibody concentration and apply a uniform immune pressure to all antibodies (see Methods).

We first validate the RBM’s learned landscape by comparing its single-site mutational effect against experimental phenotypes. For each site *i*, we computed the RBM log-likelihood score for single-site mutations in the wild-type background:

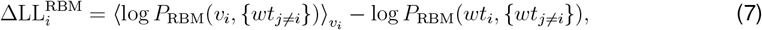

Where 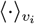 denotes the average over all amino-acid substitutions *v*_*i*_ at position *i*. This score reflects the expected effect of mutations at site *i* on protein viability, with positive values indicating mutations more favorable than the wild-type residue^39^. As shown in Figure S3, these RBM-derived scores correlate well with DMS measurements of both protein expression and ACE2 binding, indicating that the model accurately captures the viability landscape of the RBD. The model achieves a Spearman correlation of 0.60 for expression data—comparable to the DCA model of Rodriguez-Rivas *et al*.^25^ and the RBM of Huot *et al*.^32^ (both trained on SARS-CoV homologous sequences). It further reaches a Spearman correlation of 0.69 for ACE2 binding, a performance not reported in these earlier studies. This increase in predictive power is likely due to the use of synthetic homologs generated with ESM-Inverse Folding, which are structurally conditioned and more likely to retain ACE2 binding, in contrast to far SARS-CoV-2 homologs which were likely not subject to the ACE2 binding constraint. The RBM retains most of the predictive power for expression, compared to the original ESM-Inverse Folding model, while remaining equally effective at predicting binding.

We then consider escape paths of length *T* = 20, corresponding to at most 20 mutations along a trajectory, consistent with the mutational load observed in circulating variants. We project the MCMC-sampled and MF trajectories into an antigenic space defined by the top two principal components (PCs) of the antibody binding energies *γ*_*η*_ of observed sequences from the pandemic that appeared at least 100 times in Nextstrain^40^ data (Figure 3a). This allows us to directly compare model-predicted antigenic drift with real-world viral evolution. Notably, the MCMC trajectories and the MF trajectory starting from wildtype align with the direction of antigenic evolution observed in circulating variants, progressing from the wild type toward major variants of concern (VOCs) such as BA.5, BQ.1.1, and XBB. Later variants appear further along the model-predicted escape path, highlighting the model’s ability to predict the temporal and directional structure of antigenic drift. Furthermore, our framework remains robust when applied to distinct backgrounds; an MF trajectory initiated from BA.1 also converges along the same antigenic direction as the subsequently emerged variants (Figure 3a)

**Fig. 3:**
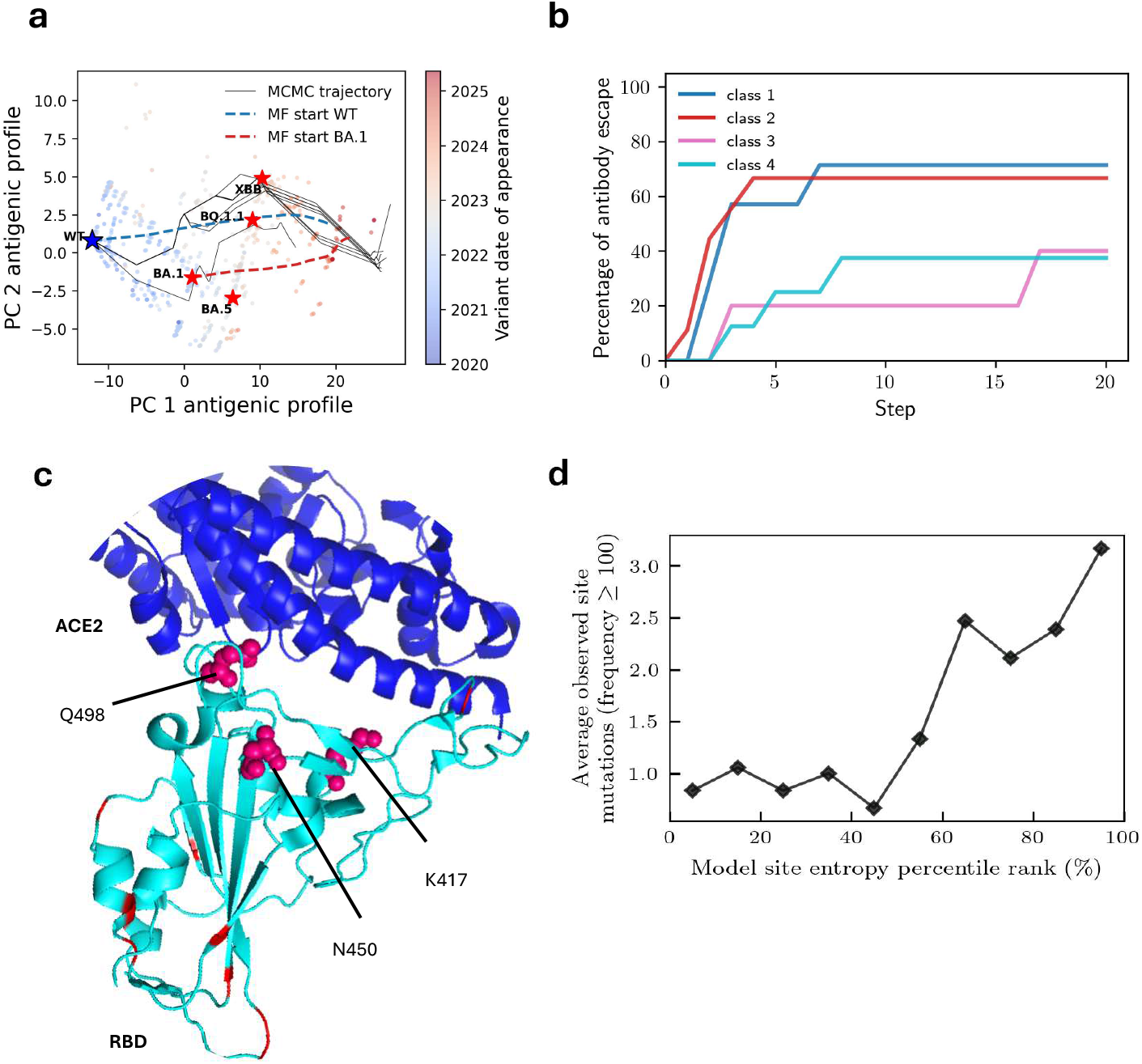
Escape paths capture mutational and antigenic features of SARS-CoV-2 variants. **(a)** Projection of evolutionary paths in antigenic space (PC1 vs. PC2 of antibody binding order parameters), comparing model predictions with pandemic surveillance data. Mean-field (MF) trajectories starting from Wildtype (WT) and BA.1, alongside MCMC trajectories from WT, are mapped against observed SARS-CoV-2 variants (colored by date of appearance). Key Variants of Concern (VOCs) are labeled. **(b)** Percentage of antibodies losing at least 90% binding along the MF trajectory, by antibody class. **(c)** Structural mapping of mutated sites in MF trajectory on the RBD (light blue) bound to ACE2 (dark blue). Red residues: mutations in the final MF sequence; Pink spheres: predicted sites also mutated in VOCs. **(d)** Comparison of site entropy from MF simulations against the number of unique mutations observed in pandemic (frequency ≥100). Sites are grouped into entropy deciles; residues in the highest entropy deciles align with sites showing extensive natural variation.

Furthermore, we validate our method by comparing binding energies of generated variants across 29 antibodies (Figure S5) along mutational paths obtained by MF predictions or MCMC simulations. The MF predictions showed strong agreement with MCMC simulations, particularly as the escape progressed (Pearson coefficients between 0.79 and 0.97). In particular, simulations result in a rapid loss of binding for Class 1 and Class 2 antibodies (Figure 3b); this is consistent with prior studies demonstrating that antibodies targeting the ACE2 receptor-binding motif—typically Classes 1 and 2—often exhibit limited breadth^15,32^.

By selecting the most probable sequence at each step of the MF trajectory (see Methods), we obtain an optimal path that leads to the escape of most considered antibodies within a few mutations. In particular, among the 13 mutations identified, we find 3 sites (450, 417, 498) that were mutated in variants of concern and are on the surface of the RBD (Figure 3c).

A key advantage of MF is that it enables the quantification of the number of viable immune escape paths through the computation of a path entropy (see Methods). This entropy reflects the diversity of trajectories consistent with both structural and immune constraints, and is reduced not only by limiting the number of mutations per step, but also by the functional constraints encoded in the RBM, which restrict the set of viable trajectories. In practice, the number of escape paths shrinks from *Q*^*NT*^ ∼10^4631^ in the unconstrained model to 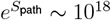. Notably, the continuity constraint of one mutation per step reduces entropy far more strongly than the RBM fields alone (site-independent model).

**Table 1:** Number of paths under different model constraints.

| Constraints | none | site-independent field | 1 mutation per step | full model |
| --- | --- | --- | --- | --- |
| Number of paths | $10^{4631}$ | $10^{1985}$ | $10^{71}$ | $10^{18}$ |

A small number of viable escape paths suggests that only specific sites can facilitate immune evasion without sacrificing protein viability. We find in Figure 3d that the site entropy calculated at the end of the MF trajectory accurately predicts the diversity of mutations observed during the pandemic (specifically those with ≥ 100 occurrences).

Altogether, these results demonstrate that our MF approximation identifies key mutational targets and offers an interpretable and efficient way to characterize dominant escape trajectories.

### Effect of cocktails

Our MF framework enables us to quantify how different antibody combinations shape viral escape dynamics and to understand why some cocktails offer greater short-term protection (evaluated here over a short trajectory of T = 10 steps). We examined the evolution of the escape probability *P*_escape_ along the MF trajectory under four different conditions: selection by individual antibodies (COV2-2196 or REGN10987) in Figures 4a–b and selection by cocktails pairing each with REGN10933 in Figures 4c–d. For each case, we identified the first point along the trajectory where the fraction of sequences with *P*_escape_ ≥ 50% is reached, providing a quantitative measure of escape speed.

**Fig. 4:**
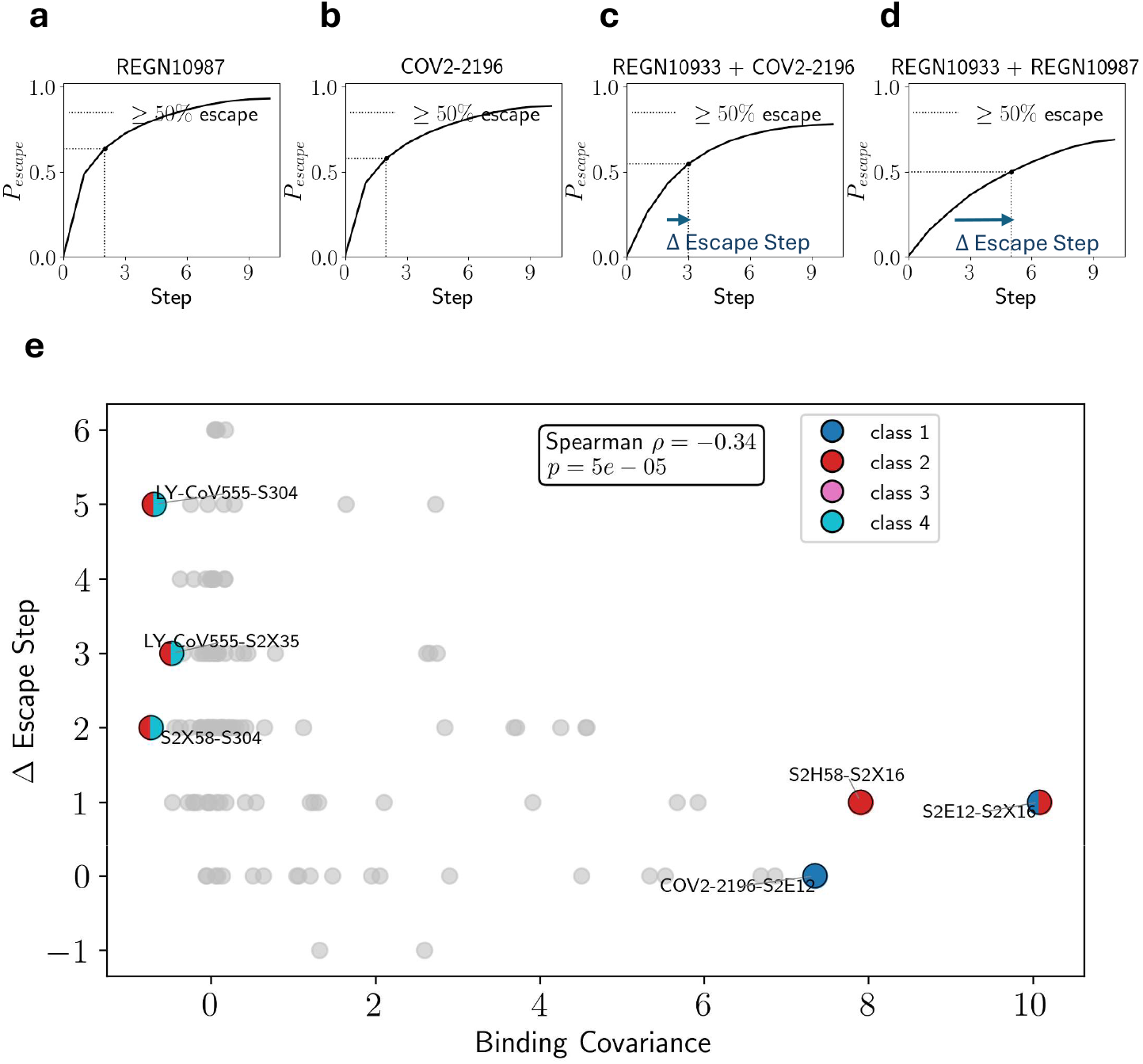
Mean field trajectories reveal resistance of antibody cocktails to viral escape. **(a–d)** Evolution of the escape probability *P*_escape_ along mean-field trajectories under different immune pressures. The virus is exposed to individual antibodies (COV2-2196 or REGN10987) and cocktails pairing each with REGN10933. The vertical dotted line indicates the step at which at least 50% escape is achieved. **(e)** Relationship between binding covariance and the delay in escape (measured as additional steps to 50% escape compared to the best antibody alone) for the 406 possible pairs of the 29 Abs considered in this work, with Spearman coefficient indicated. This difference acts as a proxy for the added evolutionary barrier imposed by the combination. Pairs with lowest/highest covariance are annotated with color indicating antibody classes.

The results show that the REGN10933 + REGN10987 cocktail substantially delays escape compared to REGN10987 alone, coherent with Baum et al.^41^, while the COV2-2196 + REGN10933 cocktail provides only marginal additional protection over COV2-2196. This suggests that not all cocktails are equally effective at slowing escape.

To understand these differences and generalize to cocktails involving all the pairs of the 29 antibodies considered in this work, we examined the correlation between escape profiles, measured as the covariance of their site-wise binding effects 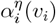 weighted by *P*_RBM_ ^32^ (Figure S6) to focus only on mutations that preserve protein viability. Analysis is restricted to pairs where the best antibody is susceptible to escape alone to allow for a defined range of escape delay values. Antibody pairs with positively correlated escape profiles tend to be evaded by the same mutations, whereas uncorrelated or anticorrelated profiles require distinct and often incompatible mutations. As shown in Figure 4e, the number of additional steps required to reach 50% escape increases when covariance between the two antibodies decreases (Spearman *ρ* = -0.34). In particular, we find that combinations of class 2 and class 4 antibodies are highly effective, whereas combining class 1 and class 2 antibodies fails to significantly delay escape. These findings agree with a previous study^32^ that demonstrated how decorrelated or anticorrelated cocktails lead to a lower number of viable escape mutants. Our work extends this by showing that these decorrelated or anticorrelated cocktails are harder to escape because they force the virus to follow longer, more constrained evolutionary paths through sequence space.

## DISCUSSION

Understanding how viruses adapt under immune pressure while preserving protein function is critical for anticipating antigenic evolution and designing therapies that remain effective over time. A core advance of our approach lies in combining generative sequence modeling with a statistical physics treatment of evolutionary paths^27^ to identify the sequential progression of mutations that enable viral escape while preserving protein viability. We leverage RBMs to capture complex epistatic patterns learned from protein sequences, including both local and higher-order dependencies that are critical for maintaining structural and functional integrity. The RBM accurately recovers folding properties in benchmark lattice models^35^ and enables distillation of sequence ensembles generated by powerful models like ESM-Inverse Folding^42^ into a simpler, tractable representation. We then apply a mean-field approximation to the ensemble of evolutionary trajectories, allowing us to analytically characterize dominant paths while quantifying key properties such as path entropy, and fitness cost.

This framework yields several key biological insights. First, we show that viability and immune constraints together lead to strongly convergent evolution, where escape trajectories rapidly collapse onto a limited number of mutational solutions. This finding is validated in both lattice protein models and the SARS-CoV-2 RBD, where we observe that mean-field generated trajectories recapitulate mutational patterns found in natural sequences. Notably, sites predicted to exhibit high mutational entropy under immune pressure correspond closely to those that have repeatedly mutated in real-world variants and most probable mutations in mean field trajectory recover observed mutated sites in pandemic.

Second, our model enables prospective forecasting of antibody escape from any variant and, crucially, incorporates evolutionary time, unlike previous approaches focused on viral variants generation^21,43^or mutational effects prediction^21–23^. By projecting trajectories into antigenic space, we find that model-predicted paths align with the direction of observed antigenic drift in SARS-CoV-2 evolution.

Third, we provide a mechanistic explanation for the effectiveness of antibody cocktails. We show that cocktails composed of antibodies with decorrelated and anti-correlated escape profiles force the virus to traverse longer paths to achieve immune evasion. These findings are not only consistent with previous empirical results^41,44,45^, but also yield a quantitative framework for optimizing cocktail combinations.

Although our framework elucidates the functional and immune constraints that shape escape funnels, it assumes a static antibody landscape and does not model the temporal evolution of the immune response. This abstraction facilitates the identification of rapid, viable escape trajectories against defined antibody sets, but excludes dynamic factors such as shifting repertoires or post-boost affinity maturation^32,46^. Therefore, rather than serving as a global epidemiological forecaster of strain dominance^2,20^, our framework functions as a mechanistic tool for mapping the accessible escape pathways driven by specific antibody pressures.

More broadly, our framework could be virus-agnostic: combining RBM-based generative modeling with statistical mechanics of constrained path ensembles makes it transferable to other rapidly evolving biological threats. RBMs and related probabilistic models have already been used to capture epistatic interactions in HIV^47^ and influenza hemagglutinin^48^ evolution. Ultimately, this model represents a significant step towards anticipating the duration over which an antibody therapy can remain effective. By mapping the number of mutations and the fitness costs required for escape, it provides a mechanistic basis for evaluating the evolutionary barriers pathogens must overcome to bypass specific therapeutic pressures.

## METHODS

### Fitness model of lattice sequences

The folding probability of a sequence **v** into a given structure *S* is determined by the energetic contributions of residue-residue interactions within the folded configuration^33,49^. These interactions occur between amino acid pairs that are spatially adjacent in the structure but not sequentially contiguous. The geometry of the structure is encoded by a contact map 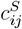, where 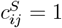 indicates that residues *i* and *j* are in contact in structure *S*, and 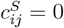 otherwise.

The total energy of a sequence **v** in a particular structure *S* is computed as:

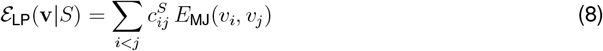

where *E*_MJ_(*v*_*i*_, *v*_*j*_) denotes the Miyazawa-Jernigan interaction energy^50^ between amino acids *v*_*i*_ and *v*_*j*_, reflecting their physico-chemical compatibility.

The probability that sequence **v** adopts structure *S* is then given by a Boltzmann distribution:

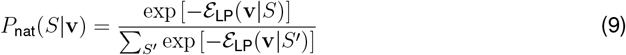

where the denominator sums over all possible compact, self-avoiding conformations *S*^*′*^ in the lattice. In this formulation, *P*_nat_ represents the ground-truth thermodynamic probability that sequence **v** adopts a given structure, based on its folding energy *ε*_LP_. However, computing this distribution exactly requires summing over all compact, self-avoiding conformations—a combinatorially expensive task for realistic protein spaces.

In our work, we therefore approximate *P*_nat_ with *P*_*RBM*_ coming from an RBM trained on sequences known to fold into the desired structure, following the procedure of Jacquin et al.^35^. Precisely, these sequences were sampled from a low temperature MC sampling using −*β* log *P*_*nat*_( | *S*_*nat*_ with −*β* = 10^2^ as effective energy.

### ESMIF distillation

For the SARS-CoV-2 RBD, we generated an artificial multiple sequence alignment (MSA) of 1,000 sequences using ESM-Inverse Folding at temperature 1, conditioned on the full RBD–ACE2 complex structure from PDB entry 6M0J. Conditioning on ACE2 binding ensures that the sampled sequences are compatible with receptor interaction, in contrast to prepandemic coronaviridae or animal SARS-related sequences that may diverge functionally. The RBM was then trained using the persistent contrastive divergence algorithm to approximate this distribution, thereby capturing structural constraints relevant for ACE2 recognition.

### Antibody binding

To model effect of single mutations on binding to RBD, we use deep mutational scans : data provided in bloomlab repository, that integrates data from several previous studies^9–15^.

We then define the binding energy contribution of amino acid *v*_*i*_ at site i to antibody *η* : 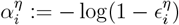, where

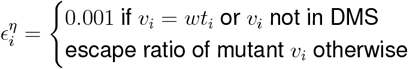

We assume that all wildtype (wt) amino acids, as well as single mutants missing from DMS, do not provide escape (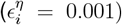). Value of 0.001 was taken as it was close to minimum measurable escape ratio among all single variants in DMS. Note that since 0 ≤ *ϵ <* 1, binding energy contributions 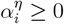.

Inspiring from Greaney et al.^38^, we then define

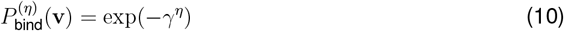

where 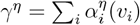 is binding energy .

In previous studies^19,32^, binding probability to an antibody with concentration *C* and dissociation constant *K*_*d*_ was typically modeled as *P*_bind_(**v**) = *C/*(*C* + *K*_*d*_(**v**)). In the low-concentration regime (*C*≪ *K*_*d*_(**v**)), *P*_bind_(**v**) ≈ *C/K*_*d*_(**v**), making the binding energy *γ*^*η*^ equal to log *K*_*d*_(**v**) used in previous approaches (up to an additive constant).

### Pandemic sequences

We downloaded SARS-CoV-2 sequence data from ([Nextstrain](https://nextstrain.org/pathogens/files?filter=ncov))^40^ on August 27, 2025. All site numbering and genome structure descriptions utilize the Wuhan-Hu-1/2019 (Genbank MN908947) strain as the reference. We used the complete dataset, containing 9’369’573 sequences, to enumerate RBD amino-acid mutation counts and analyze their appearance frequencies. From this dataset, the model is applied to sequences spanning residues S349 to G526.

### Path sampling with MCMC

We sample mutation paths under the target distribution *P*_path_. Paths start from a fixed wildtype *v*_0_ and are updated by single–mutation Metropolis–Hastings moves. Proposed updates depend on the Hamming distance between neighboring states: if neighbors coincide, a new mutation is drawn randomly; if they differ by one residue, the update is restricted to that site; and if they differ by two residues, the update flips to the only compatible intermediate. Each proposal is accepted with probability ensuring detailed balance, so that the algorithm converges to the target distribution *P*_path_ (see SI, MCMC algorithm for path sampling).

### Mean field

We extend the framework of Mauri et al.^27,28^, which formulated mean–field dynamics of mutational paths under folding constraints, in two key ways. First, we incorporate antibody escape by introducing binding–energy order parameters that act in concurrence with folding pressure, thereby capturing the trade–off between structural stability and immune evasion. Second, we free the end of the trajectories: unlike transition–path formulations that assume a fixed evolutionary destination, our paths are open-ended and evolve solely under the joint action of folding and immune constraints.

The overall probability distribution combines three potentials, coming from RBM, antibody escape and transition factors. First, the RBM likelihood reads:

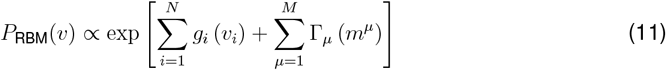

(see SI, RBM-based sequence modeling). Second, the antibody escape likelihood is given by

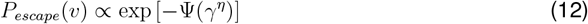

where the escape potential is

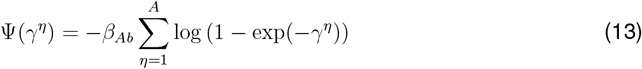

and *A* is the number of antibodies considered. *β*_*Ab*_ reflects the magnitude of immune pressure and balances the trade-off between viability and antibody escape. We assign *β*_*Ab*_ values of 10, 5, and 1 for lattice proteins, the SARS-CoV-2 RBD pandemic trajectory, and the SARS-CoV-2 RBD response to antibody cocktails, respectively. Third, to favor a low Hamming distance *d* between successive sequences along the paths, we write the transition factor in eqn. (1) as

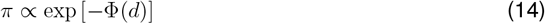

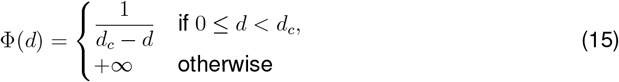

is a repulsive potential with a hard wall in *d*_*c*_ equal to the maximum number of mutations per step along the path.

Lastly, the path free energy is given by

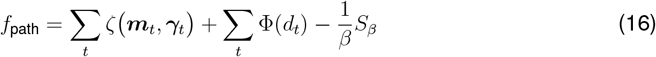

where

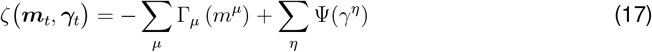

captures the tradeoff between protein viability and immune escape. The entropy term *S*_*β*_ captures the diversity of sequence realizations with the same order parameters. The mean-field trajectory is obtained by minimizing *f*_path_ with respect to the order parameters, subject to initial conditions fixed at one given sequence, such as the wildtype (see SI, Mean field).

### Most probable sequence in mean-field trajectory

In the mean-field approximation, the marginal probability 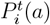 of observing amino acid a at position i and time t is derived from the derivative of the partition function with respect to the local fields.

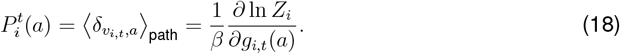

We construct the most probable sequence at each time step by selecting the amino acid a that maximizes this marginal probability at every position i.

### Site entropy

To quantify the mutational variability at each site i of the RBD, we compute the Shannon entropy:

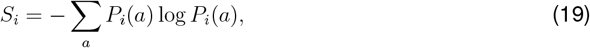

where *P*_*i*_(*a*) is the marginal probability of observing amino acid a at position i. High entropy indicates that the site tolerates diverse mutations, while low entropy reflects evolutionary constraints or functional importance.

### Path Entropy

To build intuition, we consider limiting cases for the path entropy. When paths are constrained to a single mutation per step and no external fields are applied, the entropy is given by *S*_path_ = *T* log[*N* (*Q* − 1)]. In the case where continuity is removed and no field is present, the entropy becomes *S*_path_ = *TN* log *Q*. If the field is site-independent and continuity is still removed, the entropy takes the form *S*_path_ = *TN* · ⟨*S*_site_⟩, where ⟨*S*_site_⟩ denotes the average entropy across sites induced by the field. Using the full model, path entropy (see SI, Path Entropy) can be computed as:

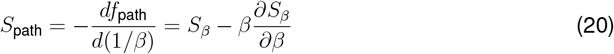

### Correlation of antibody escape profiles

To quantify the similarity between antibody escape profiles, we computed the covariance between the binding energy of antibodies identified early in the pandemic^15^. For a given antibody *η*, the binding energy, defined as *γ*^*η*^(*v*) was evaluated over sequences *v* sampled from *P*_RBM_, thereby biasing the analysis toward structurally viable variants.

To quantify antibody synergy, we compute the covariance between dissociation constants associated with antibodies 1 and 2^32^:

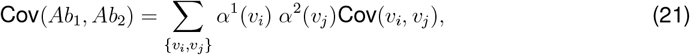

where Cov represents the covariance of amino-acid occurrences when sampling sequences from the distribution *P*_*RBM*_ . Sampling sequences from the RBM distribution ensures that escape profile correlations capture mutational accessibility under stability constraints, rather than relying solely on epitope comparison as done in previous studies^13,38^.

## RESOURCE AVAILABILITY

### Lead contact

Requests for further information and resources should be directed to and will be fulfilled by the lead contact, Simona Cocco.

### Data and code availability

The code used in this study is available at https://github.com/m-huot/ESCAPE_PATHS.

## ACKNOWLEDGMENTS

SC is grateful to G. Parisi for having suggested the application of transition paths to SARSCoV-2 evolution. We acknowledge funding from the Agence Nationale de la Recherche ( ANR ProDiGen, AAP2024 CE45 to S.C. and R.M.) and NIH R35GM139571 to E.S. The content is solely the responsibility of the authors and does not necessarily represent the official views of the National Institutes of Health.

## AUTHOR CONTRIBUTIONS

M.H., D.W., E.S., R.M., and S.C. conceived the project, supervised the research, and wrote the manuscript. M.H. coded the computational model and performed data analysis.

## SUPPLEMENTAL INFORMATION INDEX

### Supplementary methods

Supplementary figures and legends, included after main figures.

## Supplementary Materials for Constrained Evolutionary Funnels Shape Viral Immune Escape

Huot et al.

### RBM-based sequence modeling

We model the probability distribution over receptor-binding domain (RBD) sequences using *Restricted Boltzmann Machines* (RBMs), using the ([PGM package](https://github.com/jertubiana/PGM))^31^. This model is trained on homologous sequences to capture essential evolutionary, structural, and functional constraints on the RBD.

An RBM defines a joint probability distribution between protein sequences and latent variables using a bipartite graphical model. The visible layer corresponds to the amino acid sequence *v* = (*v*_1_, *v*_2_, …, *v*_*N*_ ), with each *v*_*i*_ taking one of 21 values (20 amino acids plus a gap). The hidden layer comprises real-valued units *h* = (*h*_1_, *h*_2_, …, *h*_*M*_ ) that represent learned features across the sequence.

The joint distribution over visible and hidden units is given by:

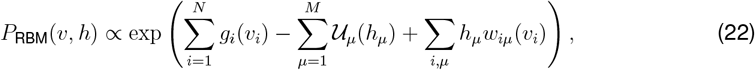

where *g*_*i*_(*v*_*i*_) and 𝒰_*μ*_(*h*_*μ*_) are local potentials (biases) on the visible and hidden units, and *w*_*iμ*_(*v*_*i*_) are interaction weights between them.

The hidden unit potential 𝒰_*μ*_(*h*) follows a double rectified linear unit (dReLU) form:

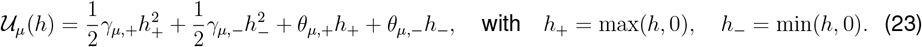

Model parameters (*γ*_*μ*,+_, *γ*_*μ,−*_, *θ*_*μ*,+_, *θ*_*μ,−*_) are estimated by maximizing the likelihood of sequences in the multiple sequence alignment (MSA), i.e., optimizing the marginal distribution:

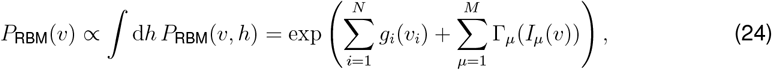

where *I*_*μ*_(*v*) =Σ_*i*_ *w*_*iμ*_(*v*_*i*_) is the input to hidden unit *μ*, and the cumulant generating function Γ_*μ*_ is defined as:

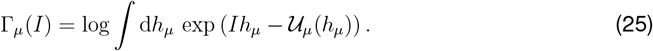

and computed following the method described in Tubiana et al.^31^.

### MCMC algorithm for path sampling

We develop a path sampling MCMC algorithm (1).

#### Algorithm 1 Metropolis–Hastings sweep over a mutation path of length *T* + 1

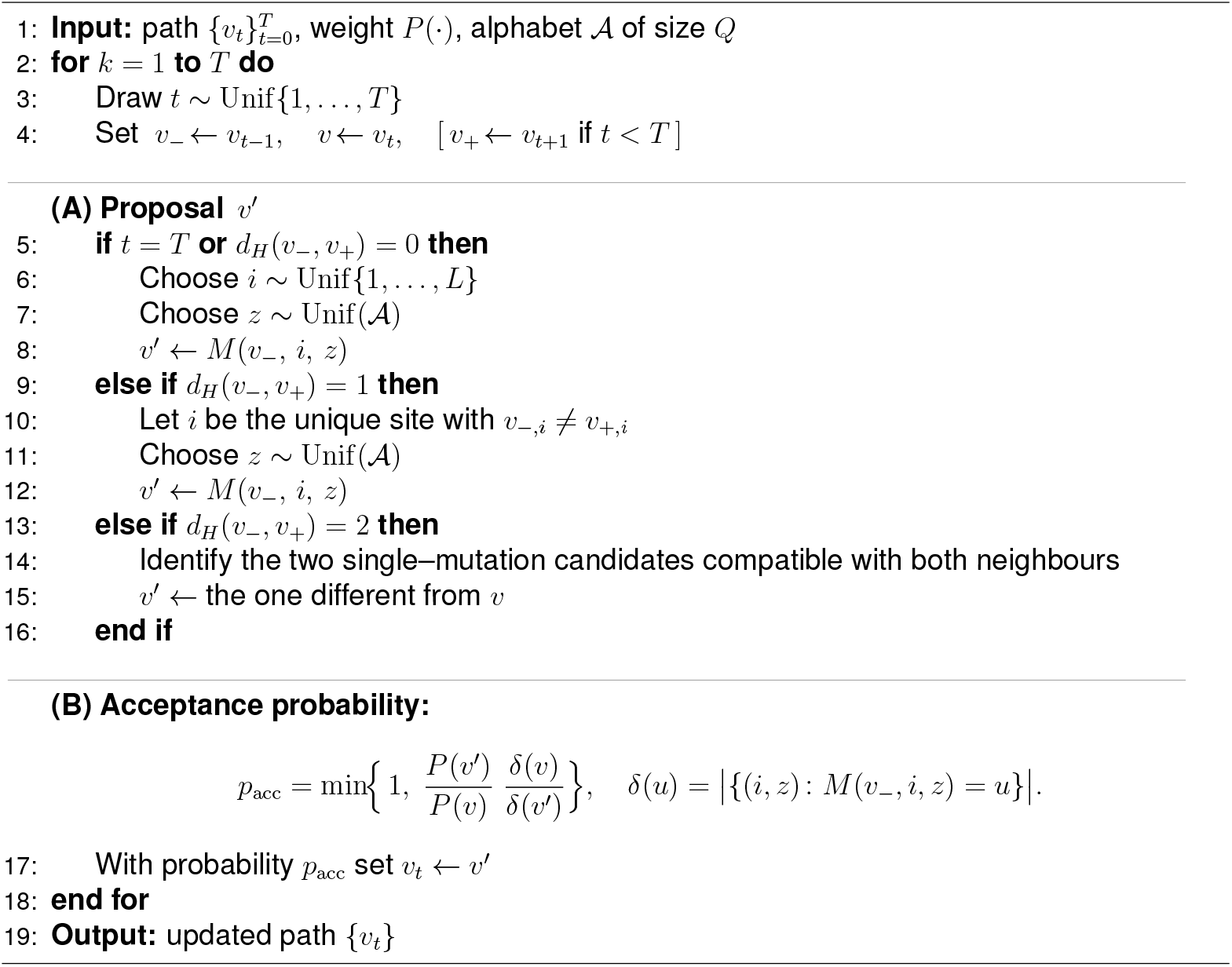

Let

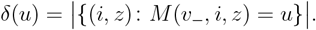

If *d*_*H*_(*v*_*−*_, *v*_+_) = 0, then

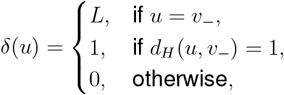

and therefore the acceptance probability

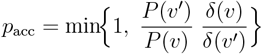

specializes to

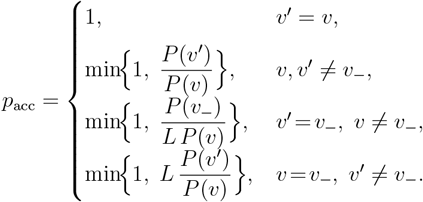

If *d*_*H*_(*v*_*−*_, *v*_+_) = 0, then every proposal and its reverse are generated in exactly one way, so *δ*(*v*) = *δ*(*v*^*′*^) = 1, and hence

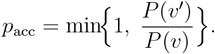

#### Proof of detailed balance

Let

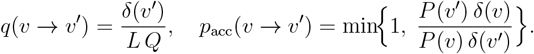

Then

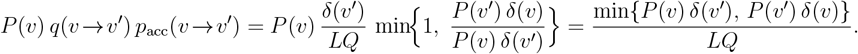

By symmetry of the minimum,

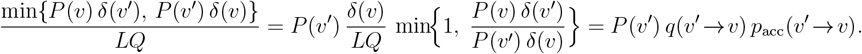

Moreover,

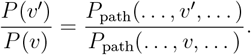

Hence 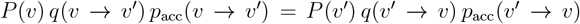 implies detailed balance for *P*_path_ as well.

**Path sampling in nucleotide space**

Path sampling in nucleotide space follows the same Metropolis–Hastings framework described above, with the key distinction that each *v*_*t*_ represents a full nucleotide sequence and each transition 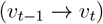 introduces at most one nucleotide mutation. The path distribution is proportional to 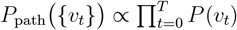 where

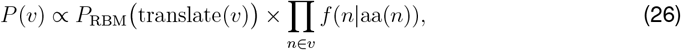

and *f* (*n* | aa) = 1*/*#codons(aa) is a uniform codon-bias term, following Di-Bari et al.^26^. Proposals introducing stop codons are automatically rejected.

### Mean field

We extend the framework developed by Mauri et al.^27,28^ for transition paths, to allow for paths that are unconstrained at their endpoints and evolve under external immune pressure. To describe the evolution of such paths, we introduce the following order parameters: the binding energy for antibody 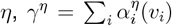 ; the input of hidden unit 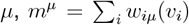 ; and the Hamming distance between successive sequences, 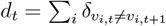.

First, RBM loglikelihood reads:

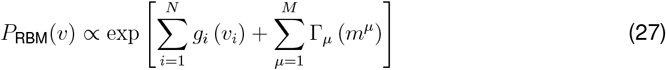

Second, antibody escape loglikelihood is given by

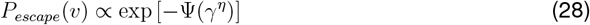

with escape potential:

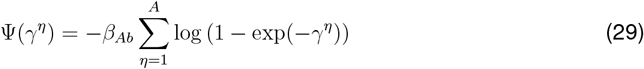

where *A* is the number of antibodies considered.

Third, to favor a low Hamming distance *d* between successive sequences along the paths, we write the transition factor loglikelihood in equation (1) as

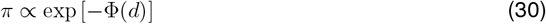

where Φ is a hard-wall repulsive potential,

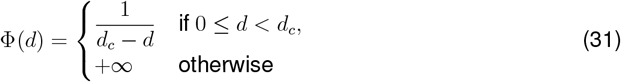

The numerical location *d*_*c*_ of the hard wall is the maximum number of mutations between two successive sequences on the path. We then define the potential 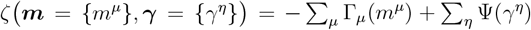 acting on the protein at each step.

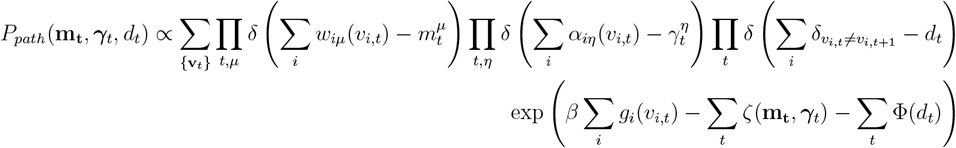

The probability of a path is given by a Boltzmann weight involving the path free energy *f*_*path*_:

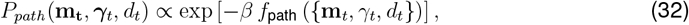

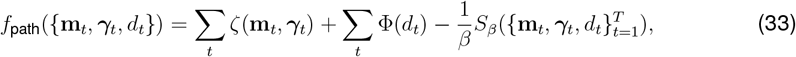

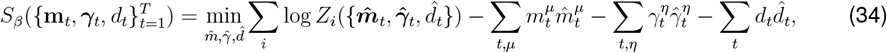

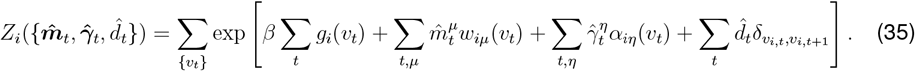

Minimizing the path free–energy density:

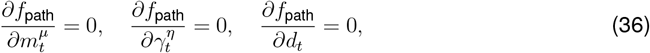

leads to the following saddle point equations for the conjugate fields:

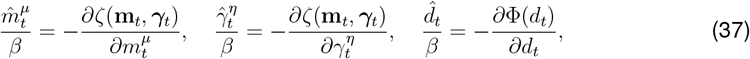

In addition, the minimization of the entropy term *S* yields the self-consistency conditions:

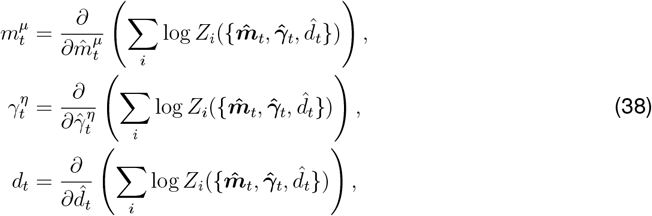

### Free energy optimisation in mean field

We first initialize (**m**_*t*_, *γ*_*t*_, *d*_*t*_) to (**m**_0_, *γ*_0_, 0), ∀*t* and apply algorithm 2.

#### Algorithm 2 Mean-Field Iteration to Compute m_*t*_, *γ*_*t*_, *d*_*t*_

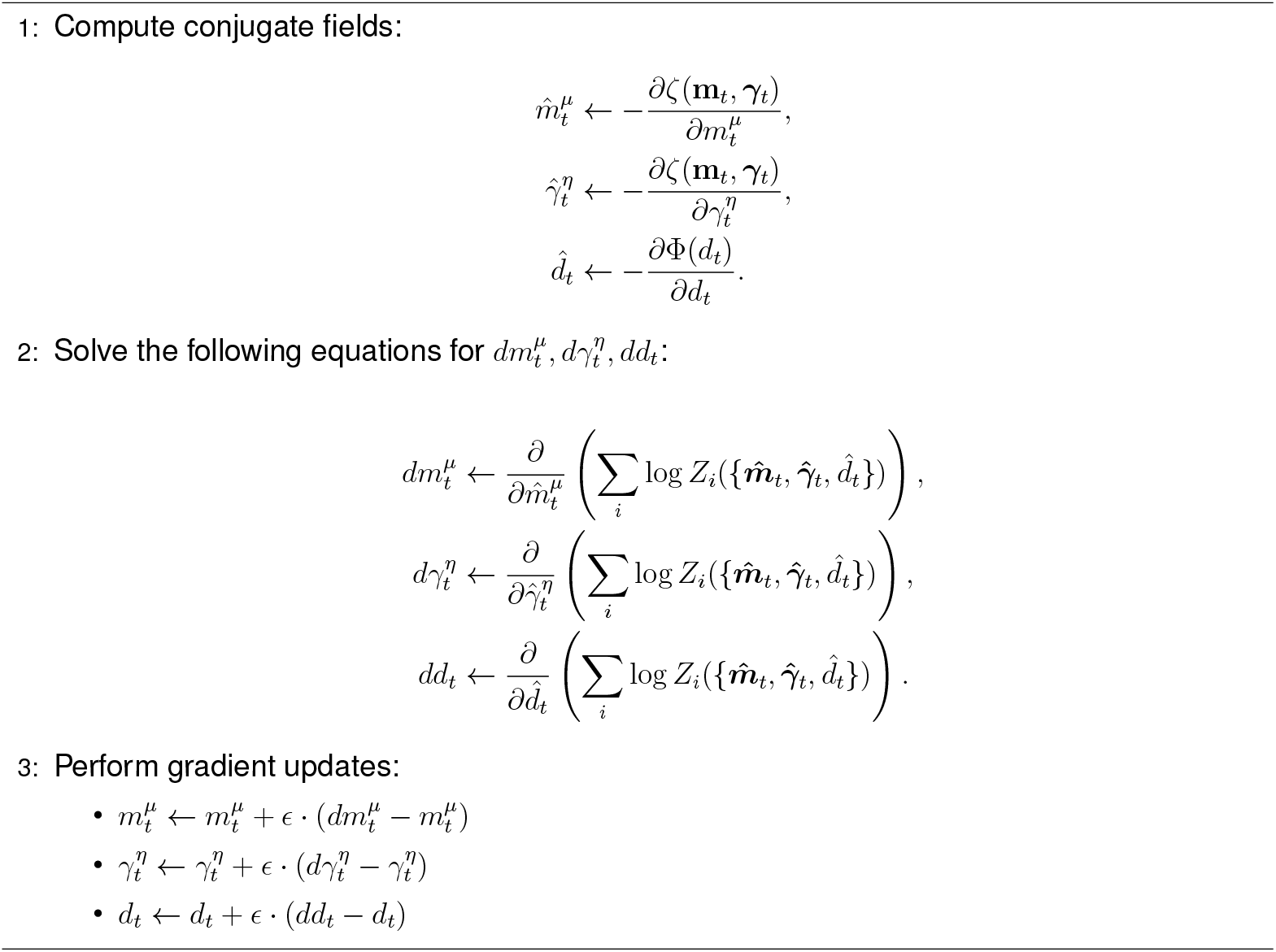

After each update of the conjugate fields, the computation of log *Z*_*i*_ is carried out using a transfer matrix method. This exploits the Markovian structure of the temporal sequence *v*_*t*_ to recursively build the partition function. .

Gradients of log *Z*_*i*_ with respect to the conjugate fields are obtained via automatic differentiation, using the PyTorch library.

### Path Entropy

To build intuition, we first consider limiting cases for the path entropy per site *S*_path_.

When paths are constrained to a single mutation per step and no external fields are applied, the entropy is given by

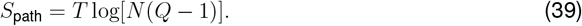

In the case where continuity is removed and no field is present, the entropy becomes

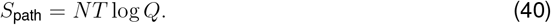

If the field is site-independent and continuity is still removed, the entropy takes the form

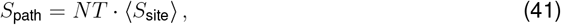

where ⟨*S*_site_⟩ denotes the average entropy across sites induced by the field.

To compute the entropy in the full mean-field model, we begin with the expression for the path free energy:

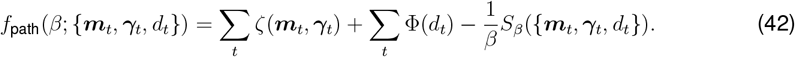

where

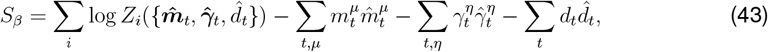

and *S*_*β*_ derivative with respect to *β* follows from the explicit dependence of *Z*_*i*_:

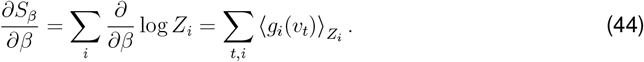

Combining all expressions, we obtain the final form for the path entropy following:

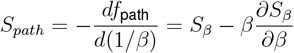

**Fig. S1:**
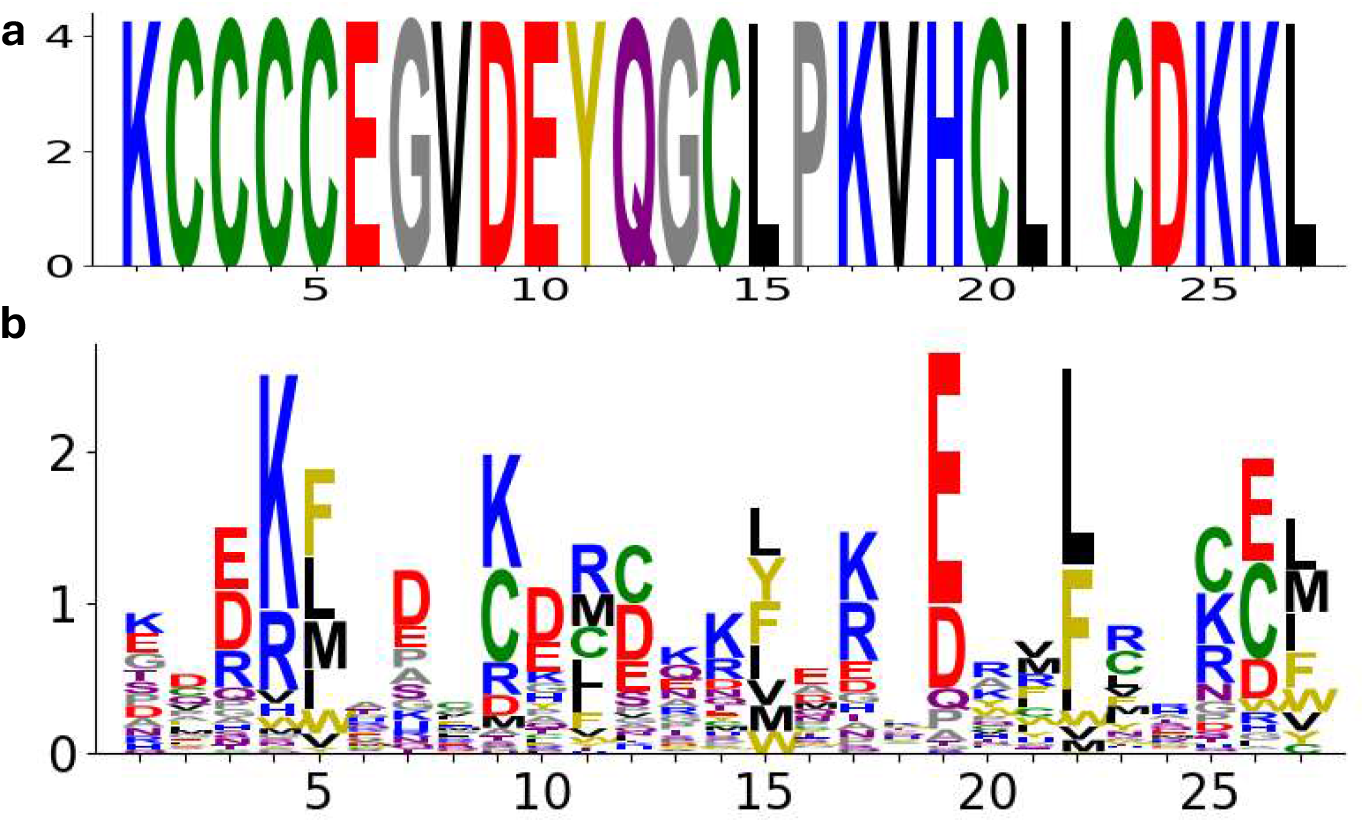
Escape paths in protein lattice model. **(a)** Wildtype sequence used as reference in the lattice model. **(b)** Sequence logo from an MSA of high-folding-probability sequences generated using the lattice model (*β* = 100). Larger letters indicate higher probability of observing a given amino acid at that site.

**Fig. S2:**
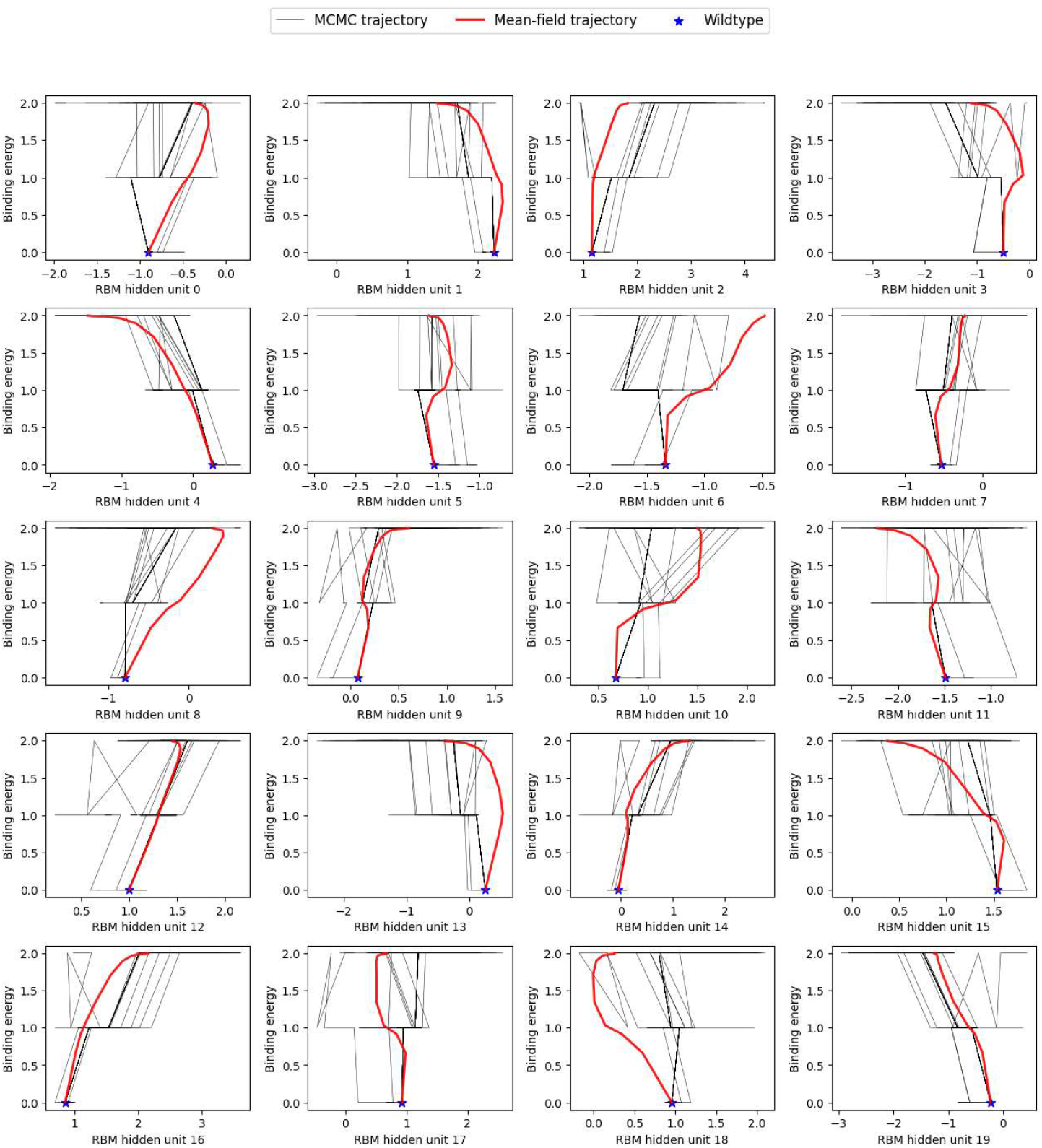
Mean field and MCMC trajectories in lattice model. Projection of mean field and MCMC trajectories in space defined by order parameters corresponding to antibody binding energy and one RBM hidden unit.

**Fig. S3:**
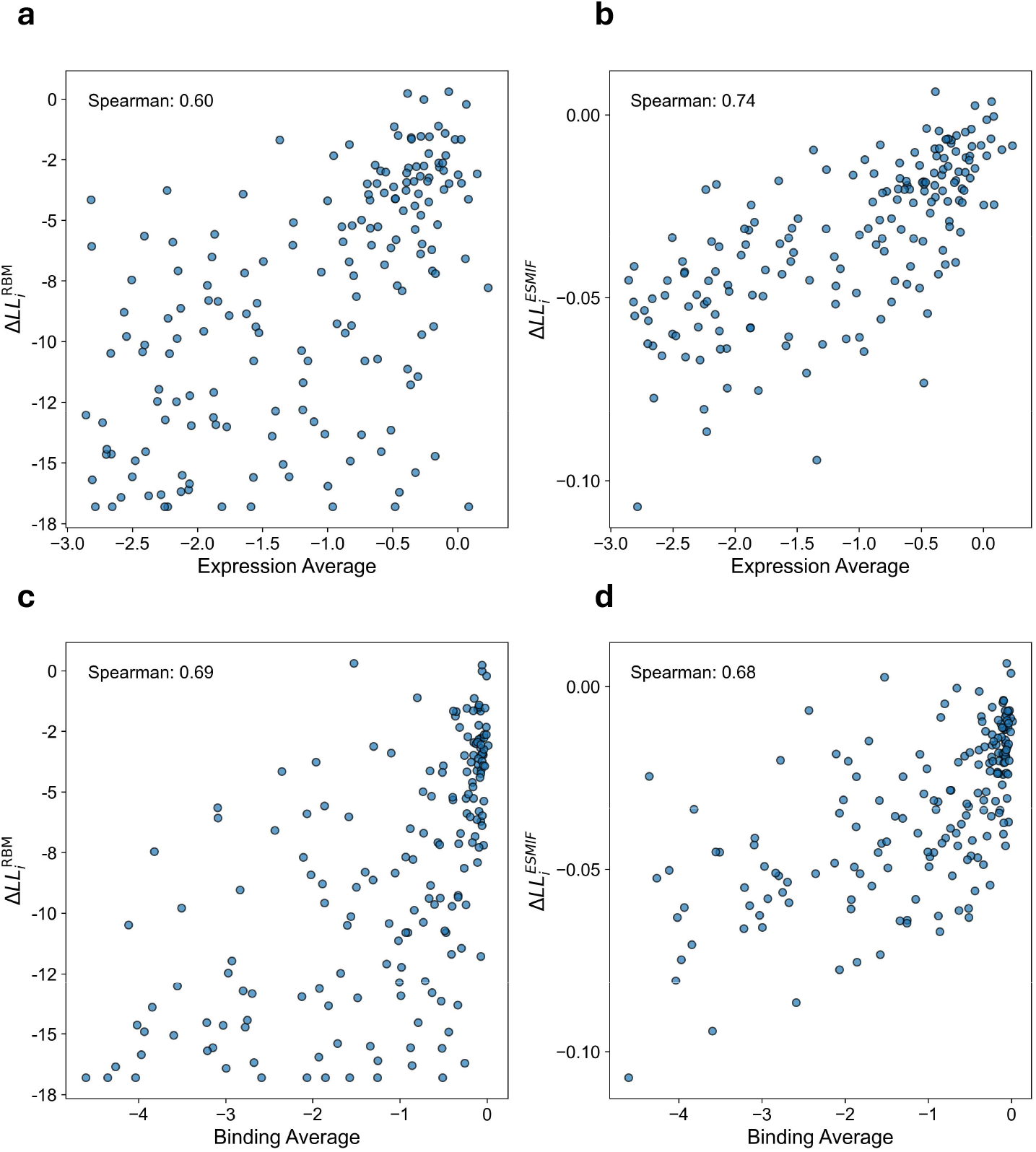
Distillation of ESMIF with RBM. **(a-b)** Comparison between experimentally measured changes in expression (Starr *et al*.), averaged per site for single mutants in the Wuhan background, and RBM/ESM-Inverse Folding–derived mutational scores (ΔLL_*i*_). **(c-d)** Same as (a-b) with experimentally measured changes in ACE2 binding (Starr *et al*.).

**Fig. S4:**
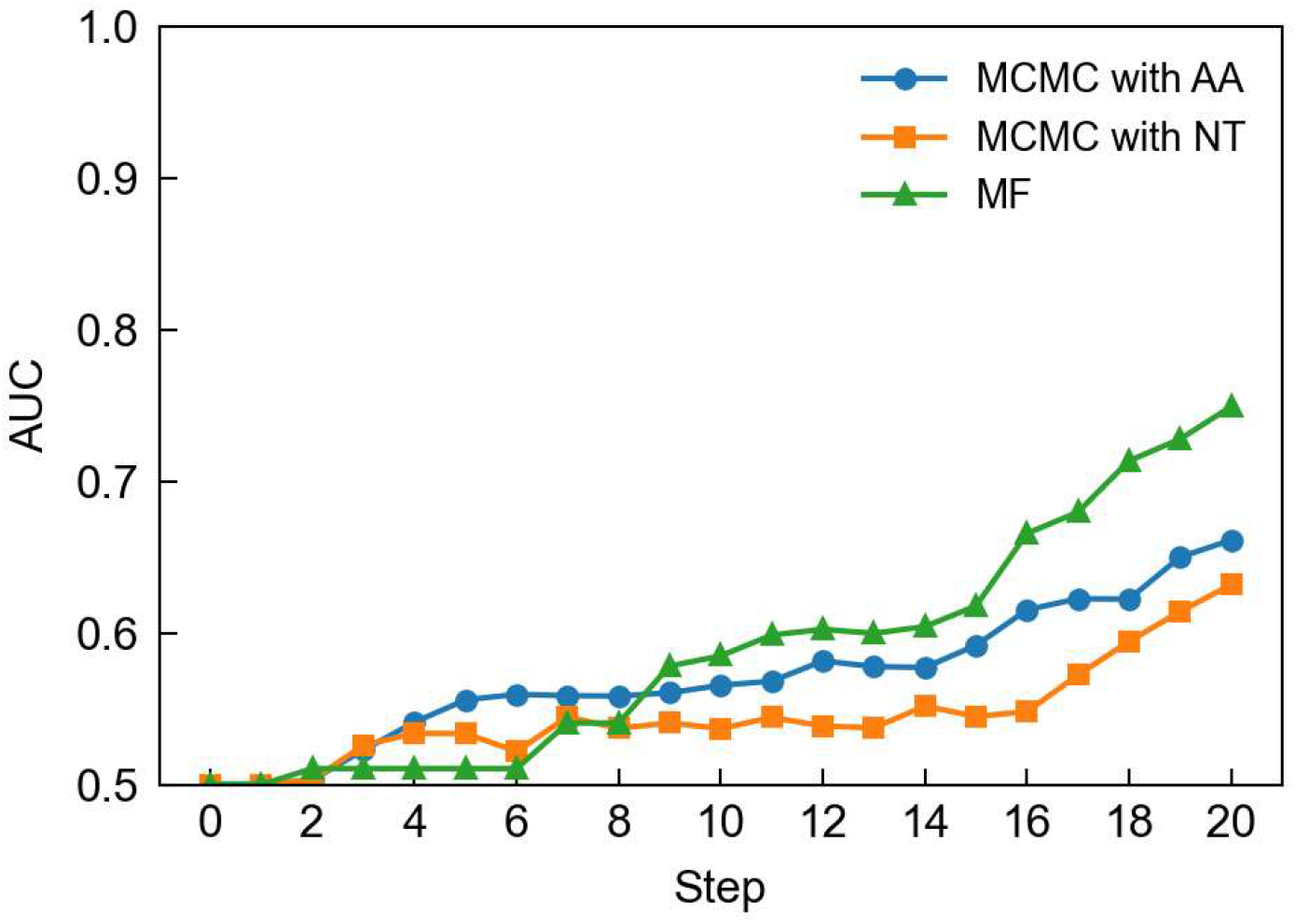
Stepwise comparison of mean field and MCMC predictions of antibody binding. Area Under the Curve (AUC) for predicting highly mutable sites in GISAID (defined as sites with ≥ 2 mutations having frequency ≥ 100). Site entropy from MF trajectory is used as a predictor and compared with site entropy from MCMC samples in amino acid (AA) or nucleotide (NT) space. AUC of 0.5 represents random predictions.

**Fig. S5.**
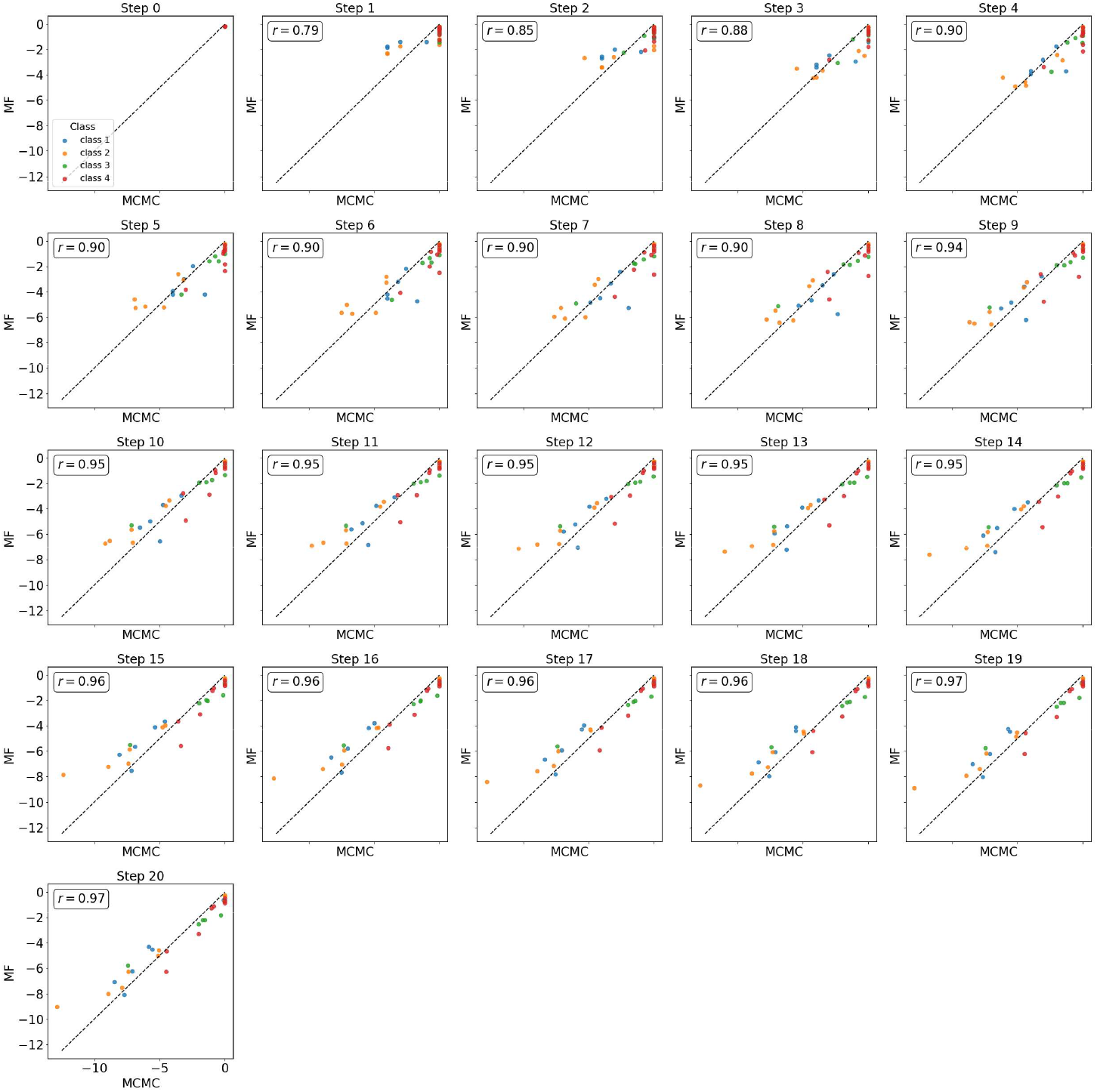
Stepwise comparison of mean field and MCMC predictions of site entropy. Correlation between antibody binding energies computed from MCMC-sampled sequences and mean field (MF) trajectory, shown for all 20 path steps. Each dot represents one antibody. Color indicates antibody class. Pearson correlation is indicated.

**Fig. S6:**
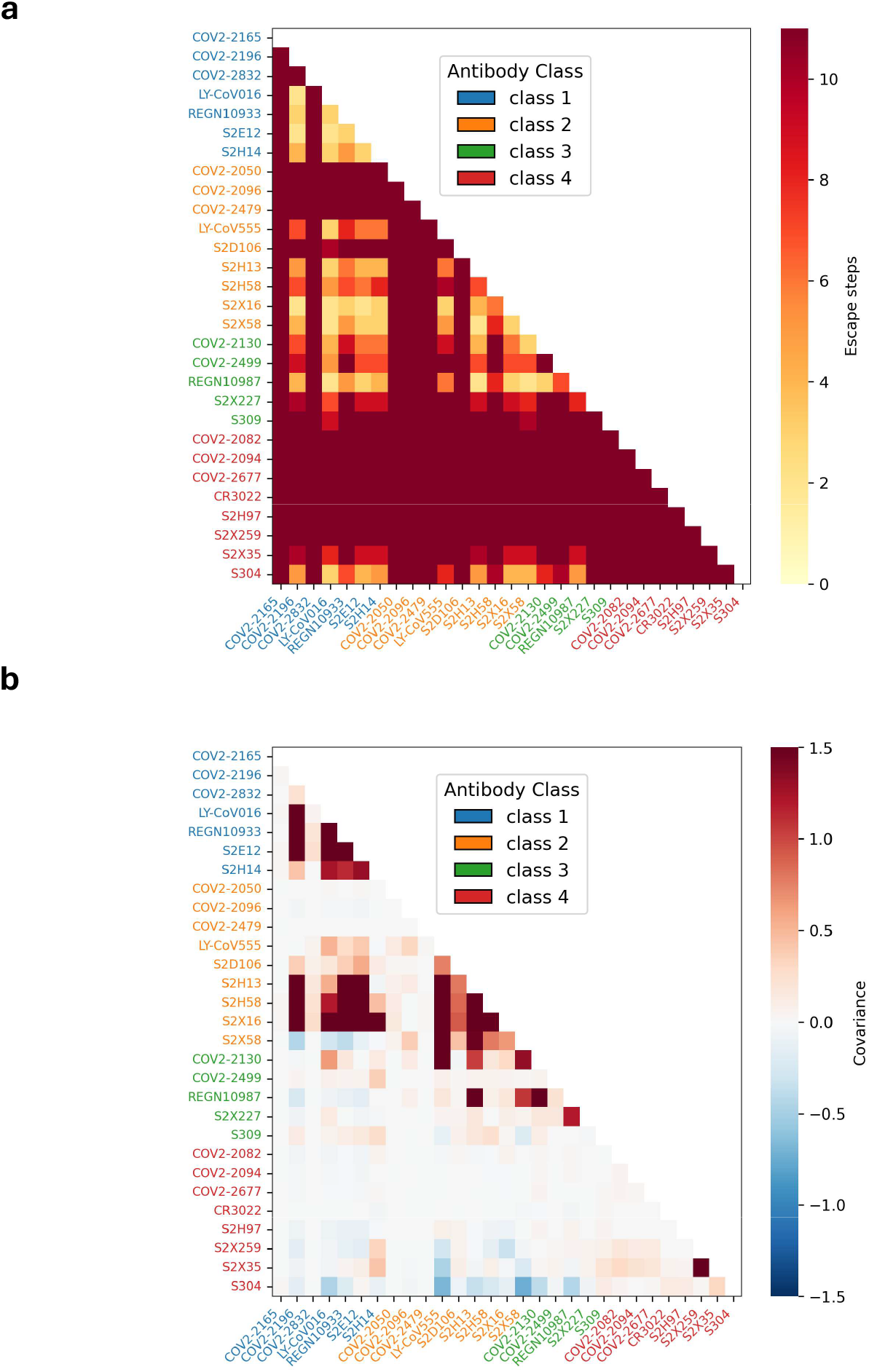
Cocktails of two antibodies. **(a)** Number of steps required for escape in cocktails of two antibodies. **(b)** Covariance between pairs of antibodies.

## References

1. Raharinirina, N.A., Gubela, N., Börnigen, D., Smith, M.R., Oh, D.Y., Budt, M., Mache, C., Schillings, C., Fuchs, S., Dürrwald, R., Wolff, T., Hölzer, M., Paraskevopoulou, S., and Von Kleist, M. (2025). SARS-CoV-2 evolution on a dynamic immune landscape. Nature 639, 196–204. URL: https://www.nature.com/articles/s41586-024-08477-8. doi: 10.1038/s41586-024-08477-8.

2. Meijers, M., Ruchnewitz, D., Eberhardt, J., Łuksza, M., and Lässig, M. (2023). Population immunity predicts evolutionary trajectories of SARS-CoV-2. Cell 186, 5151–5164.e13. URL: https://linkinghub.elsevier.com/retrieve/pii/S0092867423010760. doi: 10.1016/j.cell.2023.09.022.

3. Carabelli, A.M., Peacock, T.P., Thorne, L.G., Harvey, W.T., Hughes, J., COVID-19 Genomics UK Consortium, De Silva, T.I., Peacock, S.J., Barclay, W.S., De Silva, T.I., Towers, G.J., and Robertson, D.L. (2023). SARS-CoV-2 variant biology: immune escape, transmission and fitness. Nature Reviews Microbiology. URL: https://www.nature.com/articles/s41579-022-00841-7. doi: 10.1038/s41579-022-00841-7.

4. Nabel, K.G., Clark, S.A., Shankar, S., Pan, J., Clark, L.E., Yang, P., Coscia, A., McKay, L.G.A., Varnum, H.H., Brusic, V., Tolan, N.V., Zhou, G., Desjardins, M., Turbett, S.E., Kanjilal, S., Sherman, A.C., Dighe, A., LaRocque, R.C., Ryan, E.T., Tylek, C., Cohen-Solal, J.F., Darcy, A.T., Tavella, D., Clabbers, A., Fan, Y., Griffiths, A., Correia, I.R., Seagal, J., Baden, L.R., Charles, R.C., and Abraham, J. (2022). Structural basis for continued antibody evasion by the SARS-CoV-2 receptor binding domain. Science 375, eabl6251. URL: https://www.science.org/doi/10.1126/science.abl6251. doi: 10.1126/science.abl6251.

5. Wang, Q., Iketani, S., Li, Z., Liu, L., Guo, Y., Huang, Y., Bowen, A.D., Liu, M., Wang, M., Yu, J., Valdez, R., Lauring, A.S., Sheng, Z., Wang, H.H., Gordon, A., Liu, L., and Ho, D.D. (2023). Alarming antibody evasion properties of rising SARS-CoV-2 BQ and XBB subvariants. Cell 186, 279–286.e8. URL: https://linkinghub.elsevier.com/retrieve/pii/S0092867422015318. doi: 10.1016/j.cell.2022.12.018.

6. Cao, Y., Jian, F., Wang, J., Yu, Y., Song, W., Yisimayi, A., Wang, J., An, R., Chen, X., Zhang, N., Wang, Y., Wang, P., Zhao, L., Sun, H., Yu, L., Yang, S., Niu, X., Xiao, T., Gu, Q., Shao, F., Hao, X., Xu, Y., Jin, R., Shen, Z., Wang, Y., and Xie, X.S. (2022). Imprinted SARSCoV-2 humoral immunity induces convergent Omicron RBD evolution. Nature. URL: https://www.nature.com/articles/s41586-022-05644-7. doi: 10.1038/s41586-022-05644-7.

7. Feng, S., Reid, G.E., Clark, N.M., Harrington, A., Uprichard, S.L., and Baker, S.C. (2024). Evidence of SARS-CoV-2 convergent evolution in immunosuppressed patients treated with antiviral therapies. Virology Journal 21, 105. URL: https://virologyj.biomedcentral.com/articles/10.1186/s12985-024-02378-y. doi: 10.1186/s12985-024-02378-y.

8. Jian, F., Feng, L., Yang, S., Yu, Y., Wang, L., Song, W., Yisimayi, A., Chen, X., Xu, Y., Wang, P., Yu, L., Wang, J., Liu, L., Niu, X., Wang, J., Xiao, T., An, R., Wang, Y., Gu, Q., Shao, F., Jin, R., Shen, Z., Wang, Y., Wang, X., and Cao, Y. (2023). Convergent evolution of SARS-CoV-2 XBB lineages on receptor-binding domain 455–456 synergistically enhances antibody evasion and ACE2 binding. PLOS Pathogens 19, e1011868. URL: https://dx.plos.org/10.1371/journal.ppat.1011868. doi: 10.1371/journal.ppat.1011868.

9. Greaney, A.J., Starr, T.N., Gilchuk, P., Zost, S.J., Binshtein, E., Loes, A.N., Hilton, S.K., Huddleston, J., Eguia, R., Crawford, K.H., Dingens, A.S., Nargi, R.S., Sutton, R.E., Suryadevara, N., Rothlauf, P.W., Liu, Z., Whelan, S.P., Carnahan, R.H., Crowe, J.E., and Bloom, J.D. (2021). Complete Mapping of Mutations to the SARS-CoV-2 Spike Receptor-Binding Domain that Escape Antibody Recognition. Cell Host & Microbe 29, 44–57.e9. URL: https://linkinghub.elsevier.com/retrieve/pii/S1931312820306247. doi: 10.1016/j.chom.2020.11.007.

10. Greaney, A.J., Starr, T.N., Barnes, C.O., Weisblum, Y., Schmidt, F., Caskey, M., Gaebler, C., Cho, A., Agudelo, M., Finkin, S., Wang, Z., Poston, D., Muecksch, F., Hatziioannou, T., Bieniasz, P.D., Robbiani, D.F., Nussenzweig, M.C., Bjorkman, P.J., and Bloom, J.D. (2021). Mapping mutations to the SARS-CoV-2 RBD that escape binding by different classes of antibodies. Nature Communications 12, 4196. URL: https://www.nature.com/articles/s41467-021-24435-8. doi: 10.1038/s41467-021-24435-8.

11. Greaney, A.J., Loes, A.N., Crawford, K.H., Starr, T.N., Malone, K.D., Chu, H.Y., and Bloom, J.D. (2021). Comprehensive mapping of mutations in the SARS-CoV-2 receptor-binding domain that affect recognition by polyclonal human plasma antibodies. Cell Host & Microbe 29, 463–476.e6. URL: https://linkinghub.elsevier.com/retrieve/pii/S1931312821000822. doi: 10.1016/j.chom.2021.02.003.

12. Tortorici, M.A., Czudnochowski, N., Starr, T.N., Marzi, R., Walls, A.C., Zatta, F., Bowen, J.E., Jaconi, S., Di Iulio, J., Wang, Z., De Marco, A., Zepeda, S.K., Pinto, D., Liu, Z., Beltramello, M., Bartha, I., Housley, M.P., Lempp, F.A., Rosen, L.E., Dellota, E., Kaiser, H., Montiel-Ruiz, M., Zhou, J., Addetia, A., Guarino, B., Culap, K., Sprugasci, N., Saliba, C., Vetti, E., Giacchetto-Sasselli, I., Fregni, C.S., Abdelnabi, R., Foo, S.Y.C., Havenar-Daughton, C., Schmid, M.A., Benigni, F., Cameroni, E., Neyts, J., Telenti, A., Virgin, H.W., Whelan, S.P.J., Snell, G., Bloom, J.D., Corti, D., Veesler, D., and Pizzuto, M.S. (2021). Broad sarbecovirus neutralization by a human monoclonal antibody. Nature 597, 103–108. URL: https://www.nature.com/articles/s41586-021-03817-4. doi: 10.1038/s41586-021-03817-4.

13. Starr, T.N., Greaney, A.J., Dingens, A.S., and Bloom, J.D. (2021). Complete map of SARSCoV-2 RBD mutations that escape the monoclonal antibody LY-CoV555 and its cocktail with LY-CoV016. Cell Reports Medicine 2, 100255. URL: https://linkinghub.elsevier.com/retrieve/pii/S2666379121000719. doi: 10.1016/j.xcrm.2021.100255.

14. Starr, T.N., Greaney, A.J., Addetia, A., Hannon, W.W., Choudhary, M.C., Dingens, A.S., Li, J.Z., and Bloom, J.D. (2021). Prospective mapping of viral mutations that escape antibodies used to treat COVID-19. Science 371, 850–854. URL: https://www.science.org/doi/10.1126/science.abf9302. doi: 10.1126/science.abf9302.

15. Starr, T.N., Czudnochowski, N., Liu, Z., Zatta, F., Park, Y.J., Addetia, A., Pinto, D., Beltramello, M., Hernandez, P., Greaney, A.J., Marzi, R., Glass, W.G., Zhang, I., Dingens, A.S., Bowen, J.E., Tortorici, M.A., Walls, A.C., Wojcechowskyj, J.A., De Marco, A., Rosen, L.E., Zhou, J., Montiel-Ruiz, M., Kaiser, H., Dillen, J.R., Tucker, H., Bassi, J., Silacci-Fregni, C., Housley, M.P., Di Iulio, J., Lombardo, G., Agostini, M., Sprugasci, N., Culap, K., Jaconi, S., Meury, M., Dellota Jr, E., Abdelnabi, R., Foo, S.Y.C., Cameroni, E., Stumpf, S., Croll, T.I., Nix, J.C., Havenar-Daughton, C., Piccoli, L., Benigni, F., Neyts, J., Telenti, A., Lempp, F.A., Pizzuto, M.S., Chodera, J.D., Hebner, C.M., Virgin, H.W., Whelan, S.P.J., Veesler, D., Corti, D., Bloom, J.D., and Snell, G. (2021). SARS-CoV-2 RBD antibodies that maximize breadth and resistance to escape. Nature 597, 97–102. URL: https://www.nature.com/articles/s41586-021-03807-6. doi: 10.1038/s41586-021-03807-6.

16. Dadonaite, B., Brown, J., McMahon, T.E., Farrell, A.G., Figgins, M.D., Asarnow, D., Stewart, C., Lee, J., Logue, J., Bedford, T., Murrell, B., Chu, H.Y., Veesler, D., and Bloom, J.D. (2024). Spike deep mutational scanning helps predict success of SARS-CoV-2 clades. Nature 631, 617–626. URL: https://www.nature.com/articles/s41586-024-07636-1. doi: 10.1038/s41586-024-07636-1.

17. Yisimayi, A., Song, W., Wang, J., Jian, F., Yu, Y., Chen, X., Xu, Y., Yang, S., Niu, X., Xiao, T., Wang, J., Zhao, L., Sun, H., An, R., Zhang, N., Wang, Y., Wang, P., Yu, L., Lv, Z., Gu, Q., Shao, F., Jin, R., Shen, Z., Xie, X.S., Wang, Y., and Cao, Y. (2024). Repeated Omicron exposures override ancestral SARS-CoV-2 immune imprinting. Nature 625, 148–156. URL: https://www.nature.com/articles/s41586-023-06753-7. doi: 10.1038/s41586-023-06753-7.

18. Ito, J., Strange, A., Liu, W., Joas, G., Lytras, S., The Genotype to Phenotype Japan (G2PJapan) Consortium, Matsuno, K., Nao, N., Sawa, H., Mizuma, K., Kojima, I., Li, J., Tsubo, T., Tanaka, S., Tsuda, M., Wang, L., Oda, Y., Ferdous, Z., Shishido, K., Fukuhara, T., Tamura, T., Suzuki, R., Suzuki, S., Tsujino, S., Ito, H., Kaku, Y., Misawa, N., Plianchaisuk, A., Guo, Z., Hinay, A.A., Usui, K., Saikruang, W., Uriu, K., Kosugi, Y., Fujita, S., M. Tolentino, J.E., Chen, L., Pan, L., Li, W., Suganami, M., Chiba, M., Yoshimura, R., Yasuda, K., Iida, K., Ohsumi, N., Tanaka, S., Okumura, K., Yoshimura, K., Sadamas, K., Nagashima, M., Asakura, H., Yoshida, I., Nakagawa, S., Takaori-Kondo, A., Shirakawa, K., Nagata, K., Nomura, R., Horisawa, Y., Tashiro, Y., Kawai, Y., Takayama, K., Hashimoto, R., Deguchi, S., Watanabe, Y., Nakata, Y., Futatsusako, H., Sakamoto, A., Yasuhara, N., Hashiguchi, T., Suzuki, T., Kimura, K., Sasaki, J., Nakajima, Y., Yajima, H., Irie, T., Kawabata, R., Sasaki-Tabata, K., Ikeda, T., Nasse, H., Shimizu, R., Begum, M.M., Jonathan, M., Mugita, Y., Leong, S., Takahashi, O., Ichihara, K., Ueno, T., Motozono, C., Toyoda, M., Saito, A., Shofa, M., Shibatani, Y., Nishiuchi, T., Zahradni, J., Andrikopoulos, P., Padilla-Blanco, M., Konar, A., and Sato, K. (2025). A protein language model for exploring viral fitness landscapes. Nature Communications 16, 4236. URL: https://www.nature.com/articles/s41467-025-59422-w. doi: 10.1038/s41467-025-59422-w.

19. Wang, D., Huot, M., Mohanty, V., and Shakhnovich, E.I. (2024). Biophysical principles predict fitness of sars-cov-2 variants. Proceedings of the National Academy of Sciences 121, e2314518121. URL: https://www.pnas.org/doi/abs/10.1073/pnas.2314518121. doi: 10.1073/pnas.2314518121. arXiv: https://www.pnas.org/doi/pdf/10.1073/pnas.2314518121.

20. Maher, M.C., Bartha, I., Weaver, S., Di Iulio, J., Ferri, E., Soriaga, L., Lempp, F.A., Hie, B.L., Bryson, B., Berger, B., Robertson, D.L., Snell, G., Corti, D., Virgin, H.W., Kosakovsky Pond, S.L., and Telenti, A. (2022). Predicting the mutational drivers of future SARS-CoV-2 variants of concern. Science Translational Medicine 14, eabk3445. URL: https://www.science.org/doi/10.1126/scitranslmed.abk3445. doi: 10.1126/scitranslmed.abk3445.

21. Huot, M., Wang, D., Liu, J., and Shakhnovich, E.I. (2025). Predicting high-fitness viral protein variants with bayesian active learning and biophysics. Proceedings of the National Academy of Sciences 122, e2503742122. URL: https://www.pnas.org/doi/abs/10.1073/pnas.2503742122. doi: 10.1073/pnas.2503742122. arXiv: https://www.pnas.org/doi/pdf/10.1073/pnas.2503742122.

22. Thadani, N.N., Gurev, S., Notin, P., Youssef, N., Rollins, N.J., Ritter, D., Sander, C., Gal, Y., and Marks, D.S. (2023). Learning from prepandemic data to forecast viral escape. Nature 622, 818–825. URL: https://www.nature.com/articles/s41586-023-06617-0. doi: 10.1038/s41586-023-06617-0.

23. Wang, G., Liu, X., Wang, K., Gao, Y., Li, G., Baptista-Hon, D.T., Yang, X.H., Xue, K., Tai, W.H., Jiang, Z., Cheng, L., Fok, M., Lau, J.Y.N., Yang, S., Lu, L., Zhang, P., and Zhang, K. (2023). Deep-learning-enabled protein–protein interaction analysis for prediction of SARSCoV-2 infectivity and variant evolution. Nature Medicine 29, 2007–2018. URL: https://www.nature.com/articles/s41591-023-02483-5. doi: 10.1038/s41591-023-02483-5.

24. Dianzhuo Wang, Huot, M., Zechen Zhang, Kaiyi Jiang, Shakhnovich, E.I., and Esvelt, K.M. (2025). Without Safeguards, AI-Biology Integration Risks Accelerating Future Pandemics. URL: https://rgdoi.net/10.13140/RG.2.2.29765.15849. doi: 10.13140/RG.2.2.29765.15849. Publisher: Unpublished.

25. Rodriguez-Rivas, J., Croce, G., Muscat, M., and Weigt, M. (2022). Epistatic models predict mutable sites in SARS-CoV-2 proteins and epitopes. Proceedings of the National Academy of Sciences 119, e2113118119. URL: https://pnas.org/doi/full/10.1073/pnas.2113118119. doi: 10.1073/pnas.2113118119.

26. Bari, L.D., Bisardi, M., Cotogno, S., Weigt, M., and Zamponi, F. (2024). Emergent time scales of epistasis in protein evolution. Proceedings of the National Academy of Sciences 121, e2406807121. URL: https://www.pnas.org/doi/abs/10.1073/pnas.2406807121. doi: 10.1073/pnas.2406807121. arXiv: https://www.pnas.org/doi/pdf/10.1073/pnas.2406807121.

27. Mauri, E., Cocco, S., and Monasson, R. (2023). Mutational Paths with Sequence-Based Models of Proteins: From Sampling to Mean-Field Characterization. Physical Review Letters 130, 158402. URL: https://link.aps.org/doi/10.1103/PhysRevLett.130.158402. doi: 10.1103/PhysRevLett.130.158402.

28. Mauri, E., Cocco, S., and Monasson, R. (2023). Transition paths in Potts-like energy landscapes: General properties and application to protein sequence models. Physical Review E 108, 024141. URL: https://link.aps.org/doi/10.1103/PhysRevE.108.024141. doi: 10.1103/PhysRevE.108.024141.

29. Greenbury, S.F., Louis, A.A., and Ahnert, S.E. (2022). The structure of genotypephenotype maps makes fitness landscapes navigable. Nature Ecology & Evolution 6, 1742–1752. URL: https://www.nature.com/articles/s41559-022-01867-z. doi: 10.1038/s41559-022-01867-z.

30. Fischer, A., and Igel, C. (2014). Training restricted Boltzmann machines: An introduction. Pattern Recognition 47, 25–39. URL: https://linkinghub.elsevier.com/retrieve/pii/S0031320313002495. doi: 10.1016/j.patcog.2013.05.025.

31. Tubiana, J., Cocco, S., and Monasson, R. (2019). Learning protein constitutive motifs from sequence data. eLife 8, e39397. URL: https://elifesciences.org/articles/39397. doi: 10.7554/eLife.39397.

32. Huot, M., Rosenbaum, P., Planchais, C., Mouquet, H., Monasson, R., and Cocco, S. (2025). Generative model of sars-cov-2 variants under functional and immune pressure unveils viral escape potential and antibody resilience. bioRxiv. URL: https://www.biorxiv.org/content/early/2025/05/13/2025.05.12.653592. doi: 10.1101/2025.05.12.653592. arXiv: https://www.biorxiv.org/content/early/2025/05/13/2025.05.12.653592.full.p

33. Lau, K.F., and Dill, K.A. (1989). A lattice statistical mechanics model of the conformational and sequence spaces of proteins. Macromolecules 22, 3986–3997. URL: https://pubs.acs.org/doi/abs/10.1021/ma00200a030. doi: 10.1021/ma00200a030.

34. Loffredo, E., Vesconi, E., Razban, R., Peleg, O., Shakhnovich, E., Cocco, S., and Monasson, R. (2023). Evolutionary dynamics of a lattice dimer: a toy model for stability vs. affinity trade-offs in proteins. Journal of Physics A: Mathematical and Theoretical 56, 455002. URL: https://iopscience.iop.org/article/10.1088/1751-8121/acfddc. doi: 10.1088/1751-8121/acfddc.

35. Jacquin, H., Gilson, A., Shakhnovich, E., Cocco, S., and Monasson, R. (2016). Benchmarking Inverse Statistical Approaches for Protein Structure and Design with Exactly Solvable Models. PLOS Computational Biology 12, e1004889. URL: https://dx.plos.org/10.1371/journal.pcbi.1004889. doi: 10.1371/journal.pcbi.1004889.

36. Sautto, G., Tarr, A.W., Mancini, N., and Clementi, M. (2013). Structural and Antigenic Definition of Hepatitis C Virus E2 Glycoprotein Epitopes Targeted by Monoclonal Antibodies. Clinical and Developmental Immunology 2013, 1–12. URL: http://www.hindawi.com/journals/jir/2013/450963/. doi: 10.1155/2013/450963.

37. Lewitus, E., Li, Y., Bai, H., Pham, P., and Rolland, M. (2024). HIV-1 Gag, Pol, and Env diversified with limited adaptation since the 1980s. mBio 15, e01749–23. URL: https://journals.asm.org/doi/10.1128/mbio.01749-23. doi: 10.1128/mbio.01749-23.

38. Greaney, A.J., Starr, T.N., and Bloom, J.D. (2022). An antibody-escape estimator for mutations to the sars-cov-2 receptor-binding domain. Virus Evolution 8, veac021. URL: https://doi.org/10.1093/ve/veac021. doi: 10.1093/ve/veac021. arXiv: https://academic.oup.com/ve/article-pdf/8/1/veac021/43671163/veac021.pdf.

39. Cocco, S., Feinauer, C., Figliuzzi, M., Monasson, R., and Weigt, M. (2018). Inverse statistical physics of protein sequences: a key issues review. Reports on Progress in Physics 81, 032601. URL: https://iopscience.iop.org/article/10.1088/1361-6633/aa9965. doi: 10.1088/1361-6633/aa9965.

40. Hadfield, J., Megill, C., Bell, S.M., Huddleston, J., Potter, B., Callender, C., Sagulenko, P., Bedford, T., and Neher, R.A. (2018). Nextstrain: real-time tracking of pathogen evolution. Bioinformatics 34, 4121–4123. URL: https://academic.oup.com/bioinformatics/article/34/23/4121/5001388. doi: 10.1093/bioinformatics/bty407.

41. Baum, A., Fulton, B.O., Wloga, E., Copin, R., Pascal, K.E., Russo, V., Giordano, S., Lanza, K., Negron, N., Ni, M., Wei, Y., Atwal, G.S., Murphy, A.J., Stahl, N., Yancopoulos, G.D., and Kyratsous, C.A. (2020). Antibody cocktail to sars-cov-2 spike protein prevents rapid mutational escape seen with individual antibodies. Science 369, 1014–1018. URL: https://www.science.org/doi/abs/10.1126/science.abd0831. doi: 10.1126/science.abd0831. arXiv: https://www.science.org/doi/pdf/10.1126/science.abd0831.

42. Hsu, C., Verkuil, R., Liu, J., Lin, Z., Hie, B., Sercu, T., Lerer, A., and Rives, A. (2022). Learning inverse folding from millions of predicted structures. In K. Chaudhuri, S. Jegelka, L. Song, C. Szepesvari, G. Niu, and S. Sabato, eds. Proceedings of the 39th International Conference on Machine Learning vol. 162 of Proceedings of Machine Learning Research. PMLR pp. 8946–8970. URL: https://proceedings.mlr.press/v162/hsu22a.html.

43. Youssef, N., Gurev, S., Ghantous, F., Brock, K.P., Jaimes, J.A., Thadani, N.N., Dauphin, A., Sherman, A.C., Yurkovetskiy, L., Soto, D., Estanboulieh, R., Kotzen, B., Notin, P., Kollasch, A.W., Cohen, A.A., Dross, S.E., Erasmus, J., Fuller, D.H., Bjorkman, P.J., Lemieux, J.E., Luban, J., Seaman, M.S., and Marks, D.S. (2025). Computationally designed proteins mimic antibody immune evasion in viral evolution. Immunity 58, 1411–1421.e6. URL: https://linkinghub.elsevier.com/retrieve/pii/S1074761325001785. doi: 10.1016/j.immuni.2025.04.015.

44. Ku, Z., Xie, X., Davidson, E., Ye, X., Su, H., Menachery, V.D., Li, Y., Yuan, Z., Zhang, X., Muruato, A.E., I Escuer, A.G., Tyrell, B., Doolan, K., Doranz, B.J., Wrapp, D., Bates, P.F., McLellan, J.S., Weiss, S.R., Zhang, N., Shi, P.Y., and An, Z. (2021). Molecular determinants and mechanism for antibody cocktail preventing SARS-CoV-2 escape. Nature Communications 12, 469. URL: https://www.nature.com/articles/s41467-020-20789-7. doi: 10.1038/s41467-020-20789-7.

45. Yu, T.C., Thornton, Z.T., Hannon, W.W., DeWitt, W.S., Radford, C.E., Matsen, F.A., and Bloom, J.D. (2022). A biophysical model of viral escape from polyclonal antibodies. Virus Evolution 8, veac110. URL: https://academic.oup.com/ve/article/doi/10.1093/ve/veac110/6889254. doi: 10.1093/ve/veac110.

46. Andreano, E., Paciello, I., Pierleoni, G., Piccini, G., Abbiento, V., Antonelli, G., Pileri, P., Manganaro, N., Pantano, E., Maccari, G., Marchese, S., Donnici, L., Benincasa, L., Giglioli, G., Leonardi, M., De Santi, C., Fabbiani, M., Rancan, I., Tumbarello, M., Montagnani, F., Sala, C., Medini, D., De Francesco, R., Montomoli, E., and Rappuoli, R. (2023). B cell analyses after SARS-CoV-2 mRNA third vaccination reveals a hybrid immunity like antibody response. Nature Communications 14, 53. URL: https://www.nature.com/articles/s41467-022-35781-6. doi: 10.1038/s41467-022-35781-6.

47. Shekhar, K., Ruberman, C.F., Ferguson, A.L., Barton, J.P., Kardar, M., and Chakraborty, A.K. (2013). Spin models inferred from patient-derived viral sequence data faithfully describe HIV fitness landscapes. Physical Review E 88, 062705. URL: https://link.aps.org/doi/10.1103/PhysRevE.88.062705. doi: 10.1103/PhysRevE.88.062705.

48. Doelger, J., Kardar, M., and Chakraborty, A.K. (2022). Inferring the intrinsic mutational fitness landscape of influenzalike evolving antigens from temporally ordered sequence data. Physical Review E 105, 024401. URL: https://link.aps.org/doi/10.1103/PhysRevE.105.024401. doi: 10.1103/PhysRevE.105.024401.

49. Shakhnovich, E.I., and Gutin, A.M. (1993). Engineering of stable and fast-folding sequences of model proteins. Proceedings of the National Academy of Sciences 90, 7195–7199. URL: https://pnas.org/doi/full/10.1073/pnas.90.15.7195. doi: 10.1073/pnas.90.15.7195.

50. Miyazawa, S., and Jernigan, R.L. (1996). Residue – Residue Potentials with a Favorable Contact Pair Term and an Unfavorable High Packing Density Term, for Simulation and Threading. Journal of Molecular Biology 256, 623–644. URL: https://linkinghub.elsevier.com/retrieve/pii/S002228369690114X. doi: 10.1006/jmbi.1996.0114.

